# LRRK2 regulates innate immune responses and neuroinflammation during *Mycobacterium tuberculosis* infection

**DOI:** 10.1101/699066

**Authors:** C.G. Weindel, S.L. Bell, T.E. Huntington, K.J. Vail, R. Srinivasan, K.L. Patrick, R.O. Watson

## Abstract

Despite many connections between mutations in leucine-rich repeat kinase 2 (*LRRK2*) and susceptibility to mycobacterial infection, we know little about its function outside of the brain, where it is studied in the context of Parkinson’s Disease (PD). Here, we report that LRRK2 controls peripheral macrophages and brain-resident glial cells’ ability to respond to and express inflammatory molecules. *LRRK2* KO macrophages express elevated basal levels of type I interferons, resulting from defective purine metabolism, mitochondrial damage, and engagement of mitochondrial DNA with the cGAS DNA sensing pathway. While *LRRK2* KO mice can control *Mycobacterium tuberculosis* (Mtb) infection, they exhibit exacerbated lung inflammation and altered activation of glial cells in PD-relevant regions of the brain. These results directly implicate LRRK2 in peripheral immunity and support the “multiple-hit hypothesis” of neurodegenerative disease, whereby infection coupled with genetic defects in *LRRK2* create an immune milieu that alters activation of glial cells and may trigger PD.

## INTRODUCTION

Mutations in leucine rich repeat kinase 2 (*LRRK2*) are a major cause of familial and sporadic Parkinson’s Disease (PD), a neurodegenerative disease characterized by selective loss of dopaminergic (DA) neurons in the substantia nigra pars compacta (SNc) region of the midbrain (Cookson, 2017; Kim and Alcalay, 2017; Martin et al., 2014; Schulz et al., 2016). Despite LRRK2 having been implicated in a variety of cellular processes including cytoskeletal dynamics (Civiero et al., 2018; Kett et al., 2012; Pellegrini et al., 2017), vesicular trafficking (Herbst and Gutierrez, 2019; Sanna et al., 2012; Shi et al., 2017), calcium signaling (Bedford et al., 2016; Calì et al., 2014), and mitochondrial function (Ryan et al., 2015; Singh et al., 2019; Yue et al., 2015), its precise mechanistic contributions to triggering and/or exacerbating PD are not known.

Of all the cellular pathways affected by *LRRK2* mutations, dysregulation of mitochondrial homeostasis has emerged as a centrally important mechanism underlying PD pathogenesis and neuronal loss (Cowan et al., 2019; Panchal and Tiwari, 2019). Indeed, other PD-associated genes such as *PARK2* (Parkin), *PINK1*, and *DJ1*, all play crucial roles in mitochondrial quality control via mitophagy. LRRK2 has been implicated in mitophagy directly through interactions with the mitochondrial outer membrane protein Miro (Hsieh et al., 2016), and several lines of evidence support roles for LRRK2 in controlling mitochondrial network dynamics through interactions with the mitochondrial fission protein DRP1 (X. Wang et al., 2012). Accordingly, a number of different cell types, including fibroblasts and iPSC-derived neurons from PD patients harboring mutations in *LRRK2* exhibit increased oxidative stress and reactive oxygen species and defects in mitochondrial network integrity (Sison et al., 2018; Smith et al., 2016). Because DA neurons in the SNc have high bioenergetic needs and a unique highly-branched morphology, they are thought to be particularly sensitive to defects in mitochondrial homeostasis conferred by mutations in *LRRK2* (Surmeier et al., 2017). In spite of these well-appreciated links, LRRK2’s contribution to mitochondrial health in cells outside of the brain remains vastly understudied.

There is mounting evidence that mutations in *LRRK2* contribute to immune outcomes both in the brain and in the periphery. For example, mutations in *LRRK2* impair NF-κB signaling pathways in iPSC-derived neurons and render rats prone to progressive neuroinflammation in response to peripheral innate immune triggers (López de Maturana et al., 2016). Additionally, chemical inhibition of *LRRK2* attenuates inflammatory responses in microglia *ex vivo* (Moehle et al., 2012). Beyond these strong connections between *LRRK2* and inflammatory responses in the brain, numerous genome-wide association studies suggest that *LRRK2* is an equally important player in the peripheral immune response. Numerous single nucleotide polymorphisms (SNPs) in *LRRK2* are associated with susceptibility to mycobacterial infection (Fava et al., 2016; Marcinek et al., 2013; D. Wang et al., 2015; F.-R. Zhang et al., 2009), inflammatory colitis (Umeno et al., 2011), and Crohn’s Disease (Van Limbergen et al., 2009). Consistent with a role for LRRK2 in pathogen defense and autoimmunity, it is abundant in many immune cells (e.g. B cells, dendritic cells, monocytes, macrophages), and expression of *LRRK2* is induced in human macrophages treated with IFN-γ (Gardet et al., 2010). Loss of *LRRK2* reduces IL-1β secretion in response to *Salmonella enterica* infection in macrophages *ex vivo* (Liu et al., 2017) and enhances expression of pro-inflammatory cytokines in response to *Mycobacterium tuberculosis* (Mtb) infection (Härtlova et al., 2018). However, the precise mechanistic contributions of *LRRK2* to controlling immune responses in the periphery remain poorly understood.

Here, we provide the first evidence that LRRK2’s ability to influence inflammatory gene expression in macrophages is directly linked to its roles in maintaining mitochondrial homeostasis. Specifically, we demonstrate that depolarization of the mitochondrial network and hyper-activation of DRP1 in *LRRK2* KO macrophages leads to the release of mtDNA, engagement of the cGAS-dependent DNA sensing pathway, and abnormally high basal levels of interferon-β (type I interferon (IFN)) and interferon stimulated genes (ISGs). These high basal levels of type I IFN appear to completely reprogram *LRRK2* KO macrophages, rendering them refractory to a number of distinct innate immune stimuli, including infection with the important human lung pathogen, Mtb. While Mtb-infected *LRRK2* KO mice did not exhibit significant differences in bacterial burdens, we did observe exacerbated pathology in the lungs. Remarkably, although no bacilli were present in the brains of control (CT) or KO mice, both exhibited dramatic signs of neuroinflammation, evidenced by activation of microglia and astrocytes in several PD-relevant brain regions. Collectively, these results demonstrate that LRRK2’s role in maintaining mitochondrial homeostasis is critical for proper induction of inflammatory gene expression in both peripheral macrophages and brain-resident glial cells. Moreover, this provides strong support for the “multiple-hit hypothesis” of neurodegeneration, whereby peripheral infection coupled with specific genetic mutations may trigger or exacerbate neuronal loss.

## RESULTS

### RNA-seq analysis reveals that *LRRK2* deficiency in macrophages results in dysregulation of the type I IFN response during Mtb infection

To begin implicating *LRRK2* in the peripheral immune response, we took an unbiased approach to determine how loss of *LRRK2* impacts innate immune gene expression during Mtb infection of macrophages *ex vivo*. Briefly, we infected primary murine bone marrow-derived macrophages (BMDMs) derived from littermate heterozygous (control, CT) and knockout (KO) *LRRK2* mice with Mtb at an MOI of 10 and performed RNA-seq analysis on total RNA from uninfected and infected cells 4 h post-infection. Previous studies have identified 4 h as a key innate immune time point during Mtb infection, corresponding to the peak of transcriptional activation downstream of sensing molecules (Manzanillo et al., 2012; Watson et al., 2015; 2012). Mtb is a potent activator of type I IFN expression, thought to occur mostly through permeabilization of the Mtb-containing phagosome and release of bacterial dsDNA into the cytosol, where it is detected by DNA sensors like cGAS to activate the STING/TBK1/IRF3 axis (Collins et al., 2015; Wassermann et al., 2015; Watson et al., 2015; Wiens and Ernst, 2016).

Following analysis with CLC Genomics Workbench, we identified hundreds of genes that were differentially expressed in *LRRK2* KO vs. CT BMDMs during Mtb infection, with 192 genes significantly up- or down-regulated (179 up, 13 down) (p<0.001) (Fig. 1A, Table S1). Although a number of genes were significantly induced during infection in both genotypes, the level of induction for a number of transcripts was noticeably lower in *LRRK2* KO cells compared to CT (Fig. 1C). Canonical pathway analysis of differentially expressed genes revealed significant enrichment for genes involved in type I IFN and other related pathways (-log(p)=12.98), including activation of IRF by cytosolic pattern recognition receptors (-log(p)=11.81), RIG-I signaling (-log(p)=4.69), and autophagy (-log(p)=3.815)(Fig. 1B, S1A). Indeed, a majority of the top differentially expressed genes were well-known interferon-stimulated genes (ISGs), including *Ifit1, Mx1*, and *Isg15* (Fig. 1C). Follow-up RT-qPCR analysis confirmed that *Ifnb* and ISGs like *Mx1, Isg15*, and *Gbp7* were induced to lower levels in *LRRK2* KO macrophages compared to controls following Mtb infection (Fig. 1D). This differential response seemed to be specific for type I IFN and ISGs since the transcripts of Mtb-induced cytokines like *Tnfa* and *Il1b* reached similar levels in both genotypes (Fig. 1E).

**Figure 1.**
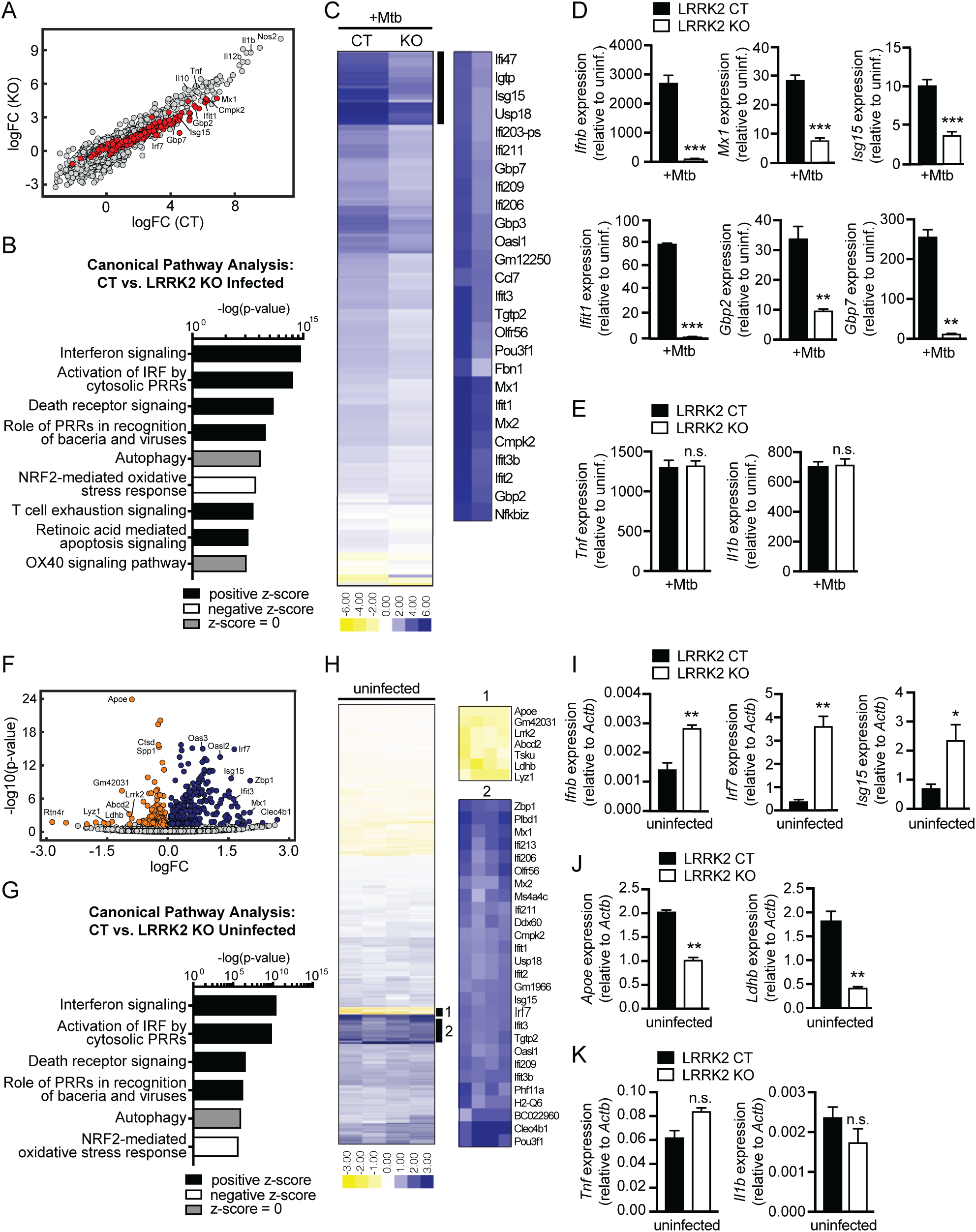
Global gene expression analysis reveals *LRRK2* KO macrophages are deficient at inducing type I IFN expression and have elevated basal type I IFN expression. **(A)** Scatter plot of genes up- and down-regulated in *LRRK2* knockout (KO) and control (CT) BMDMs 4 h post-infection with Mtb. Genes whose expression changes are significant (p<0.001; adjusted p-value is p<0.05) are highlighted in red. **(B)** IPA software analysis of cellular pathways enriched for differentially expressed genes in *LRRK2* KO vs. CT BMDMs during Mtb infection**. (C)** Heatmap of significant gene expression differences (log2fold-change) in *LRRK2* KO vs. CT BMDMs during Mtb infection. **(D)** RT-qPCR of fold-change in transcripts for type I IFN genes (*Ifnb, Mx1, Isg15*) and type II IFN genes (*Ifit1, Gbp2, Gbp7*) during Mtb infection**. (E)** RT-qPCR of NF-κB genes (*Tnfa* and *Il1b)* during Mtb infection**. (F)** Volcano plot of genes significantly upregulated (blue) or downregulated (orange) in uninfected (resting) *LRRK2* KO BMDMs**. (G)** As in (B) but comparing uninfected *LRRK2* KO vs. CT BMDMs**. (H)** As in (C) but for uninfected *LRRK2* KO vs. CT BMDMs. Zoom 1 is top downregulated genes; Zoom 2 is top upregulated genes. **(I)** RT-qPCR of type I IFN associated genes (*Ifnb, Irf7, Isg15)* normalized to *Actb* in uninfected BMDMs**. (J)** RT-qPCR of *Apoe* and *Ldhb* normalized to *Actb* in uninfected BMDMs. **(K)** As in (E) but in uninfected BMDMs. Data represented as means +/- S.E.M. *p<0.05, **p<0.01, ***p<0.005. See also Figure S1 and Table S1.

### Resting *LRRK2* KO macrophages express elevated levels of type I IFN

To begin investigating the nature of this defect in type I IFN induction, we again used CLC Genomics Workbench analysis to identify transcripts affected by loss of *LRRK2* in uninfected or “resting” macrophages. We observed higher basal expression of a number of innate immune transcripts in *LRRK2* KO BMDMs relative to control, including several type I IFN family genes (e.g. *Oas3, Irf7, Oasl2, Isg15, Zbp1*) (Fig. 1F-H). Indeed, differential gene expression analysis and unbiased canonical pathways analysis again revealed that “Interferon Signaling” was the most significantly impacted pathway in resting macrophages (-log(p)=12.95) (Fig. 1G). In fact, almost all the pathways and families of genes differentially expressed in *LRRK2* KO BMDMs in *Mtb*-infected cells were also impacted in uninfected cells (Fig. 1B and G, S1A-D). RT-qPCR analysis confirmed significantly elevated levels of type I IFN transcripts including *Ifnb, Irf7, Isg15*, and *Ifit1* in uninfected *LRRK2* KO BMDMs (Fig. 1I and S1E). Interestingly, several transcripts, including *Apoe*, a gene associated with Alzheimer’s and cardiovascular disease, had decreased expression in *LRRK2* KO BMDMs compared to controls (Fig. 1J). Transcripts like *Tnfa* and *Il1b* were expressed at similar levels in the two genotypes in resting macrophages (Fig. 1K). Importantly, the basal and induced levels of interferon and ISGs were similar between wildtype and heterozygous *LRRK2* BMDMs, validating our use of heterozygous littermates as controls in future experiments (Fig. S1F-G).

We also found increased basal expression and decreased induction of interferon and ISGs upon Mtb infection in the human monocyte cell line U937 (Fig. 2A and S2A) and in RAW 264.7 macrophages when *LRRK2* was knocked down by shRNA (Fig. S2C). *LRRK2* KO RAW 264.7 cells infected with *Mycobacterium leprae*, which like Mtb has a virulence-associated ESX-1 secretion system and induces type I IFN through cytosolic nucleic acid sensing (de Toledo-Pinto et al., 2016), had a similar defect in type I IFN induction compared to control cells (Fig. 2B and S2B). Together, these transcriptome-focused analyses revealed that *LRRK2* KO macrophages have a higher interferon signature at baseline but are unable to induce the type I IFN response to the same levels as controls when infected with Mtb or *M. leprae*.

**Figure 2.**
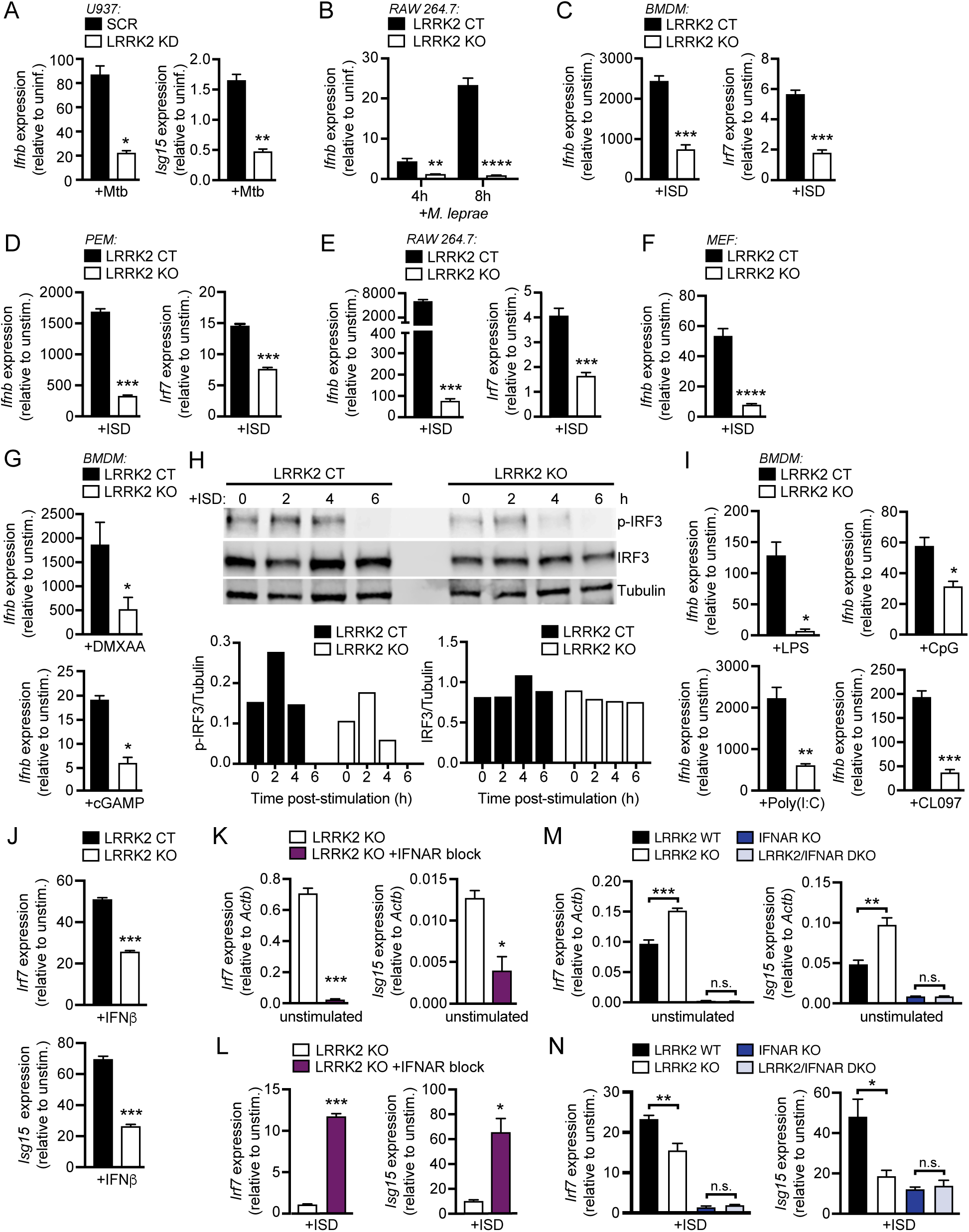
Loss of *LRRK2* contributes to type I IFN dysregulation independently of nucleic acid sensing or IFNAR signaling. **(A)** RT-qPCR of fold-change in *Ifnb* and *Isg15* expression during Mtb infection in differentiated U937 monocytes stably expressing scramble shRNA (SCR) or shRNA targeted to *LRRK2* (KD)**. (B)** RT-qPCR of fold-change in *Isg15* expression at 4 and 8 h post-infection with *M. leprae* in CT or *LRRK2* KO RAW 264.7 cells. **(C-F)** RT-qPCR of fold-change in *Ifnb* and *Irf7* expression 4 h post-transfection with 1μg/ml ISD (dsDNA) in *LRRK2* KO vs. CT (C) BMDMs, (D) peritoneal macrophages (PEMs), (E) RAW 264.7, and (F) primary mouse embryonic fibroblasts (MEFs). **(G)** RT-qPCR of fold-change in *Ifnb* expression in *LRRK2* CT or KO BMDMs following stimulation with 50 ng/ml DMXAA (2 h) or 1 μg/ml cGAMP (4 h). **(H)** Quantitative western blot of IRF3 phosphorylated at Ser396 (p-IRF3), total IRF3, and tubulin at 0, 2, 4, and 6h post-transfection with 1 μg/ml ISD in *LRRK2* KO vs. CT BMDMs. Lower graphs show quantification of p-IRF3 and IRF3 to tubulin. **(I)** As in (G) with 100 ng/ml LPS (4 h), transfection of 10 μM CpG 2395 (4 h), 1 mg/ml poly(I:C) (dsRNA) (4 h) or 1 μM CL097 (4 h). **(J)** RT-qPCR of fold-change in *Irf7* and *Isg15* expression following stimulation with 200IU IFN-β (4 h) in LRRK2 KO vs. CT BMDMs. **(K)** RT-qPCR of *Irf7* and *Isg15* expression normalized to *Actb* in *LRRK2* KO BMDMs in the presence of overnight blocking with IFN-β neutralizing antibody (1:250). **(L)** As in (J) but fold-change in *Irf7* and *Isg15* expression after transfection of 1 μg/ml ISD for 4h. **(M)** As in (K) but in BMDMs from CT, *LRRK2* KO, *IFNAR* KO, and *LRRK2*/*IFNAR* double KO double (DKO) mice. **(N)** As in (M) but fold-change in *Irf7* and *Isg15* expression after transfection of 1 μg/ml ISD for 4h. Data represented as means +/- S.E.M. *p<0.05, **p<0.01, ***p<0.005. See also Figure S2.

### *LRRK2* KO macrophages fail to induce type I IFN in response to diverse innate immune stimuli

Because both Mtb and *M. leprae* stimulate type I IFN through the cGAS/STING/TBK1 axis, we hypothesized that loss of *LRRK2* may cause defects in this pathway. To begin testing this, we stimulated a variety of *LRRK2*-deficient cells via transfection of interferon-stimulating DNA (ISD). *LRRK2* KO BMDMs failed to fully induce *Ifnb* and *Irf7* following stimulation (Fig. 2C), as did *LRRK2* KO peritoneal macrophages (PEMs) (Fig. 2D), *LRRK2* KO or KD RAW 264.7 macrophages (Fig. 2E and S2C), and *LRRK2* KO primary mouse embryonic fibroblasts (MEFs) (Fig 2F and S2D). Likewise, direct stimulation of STING with the agonist DMXAA or with transfection of the cGAS second messenger cGAMP also failed to induce type I IFN in *LRRK2* KO BMDMs (Fig. 2G), *LRRK2* KO PEMs (Fig. S2F), and *LRRK2* KO RAW 264.7 cells (Fig. S2F). Consistent with our RT-qPCR data, western blot analysis of IRF3 (Ser 395) after ISD transfection showed a significant defect in the ability of *LRRK2* KO BMDMs to respond to cytosolic DNA (Fig. 2H).

We next tested whether loss of *LRRK2* impacts the ability of cells to respond to other innate immune agonists that elicit a type I IFN response. To this end, we treated *LRRK2* KO and CT BMDMs with poly(I:C) (via transfection; to activate RNA sensing), LPS (to stimulate TRIF/IRF3 downstream of TLR4), CpG (to stimulate nucleic acid sensing via TLR9), and CL097 (to stimulate nucleic acid sensing via TLR7). In all cases, we observed a defect in the ability of *LRRK2* KO BMDMs to induce *Ifnb* (Fig. 2I). Likewise, *LRRK2* KO MEFs and PEMs also failed to induce Ifnb and Irf7 (Fig. S2G-H). *LRRK2* KO BMDMs were also defective in ISG expression (*Isg15* and *Irf7*) following treatment with recombinant bioactive IFN-β, which directly engages the interferon-alpha/beta receptor (IFNAR) (Fig. 2J). Based on these collective results, we concluded that *LRRK2* KO macrophages are reprogrammed such that they cannot properly induce type I IFN expression regardless of the innate immune stimulus received.

We next hypothesized that the elevated basal levels of type I IFN transcripts were driving the inability of *LRRK2* KO macrophages to properly induce a type I IFN response. To test this, we treated CT and KO *LRRK2* BMDMs with an IFN-β neutralizing antibody to prevent engagement of IFNAR and downstream signaling events. As predicted, this IFN-β blockade decreased basal levels of *Irf7* and *Isg15* in *LRRK2* KO cells (Fig. 2K), and remarkably, it restored the ability of *LRRK2* KO BMDMs to fully induce type I IFN expression following ISD transfection (Fig. 2L). We further tested if loss of IFNAR signaling could rescue the *LRRK2* KO phenotype by crossing *LRRK2* KO mice to *IFNAR* KO mice. In *LRRK2*/*IFNAR* double KO BMDMs, we observed a significant reduction in basal ISG levels (Fig. 2M). Upon stimulation, induction of ISGs in the *LRRK2/IFNAR* double KO BMDMs was similar to *IFNAR* KO control cells (Fig 2N). These results indicate that cytosolic nucleic acid sensing and IFNAR signaling are intact in *LRRK2* KO macrophages, but chronic elevated basal type I IFN expression renders these cells refractory to innate immune stimuli.

### Increased basal IFN in *LRRK2* KO macrophages is dependent on cytosolic DNA sensing through cGAS

Because IFN-β blockade and loss of IFNAR normalized ISG expression in *LRRK2* KO macrophages, we hypothesized that LRRK2 contributes to basal type I IFN expression upstream of the cytosolic DNA sensing pathway. To directly test the involvement of cGAS in generating elevated resting type I IFN in *LRRK2* KO macrophages, we crossed *LRRK2* KO and *cGAS* KO mice and compared type I IFN transcripts in *LRRK2/cGAS* double KO BMDMs with littermate controls. As expected, loss of *cGAS* led to lower resting *Ifnb, Isg15*, and *Irf7* expression (Fig. 3A and S3A) (Schoggins et al., 2014). Importantly, knocking out *cGAS* in a *LRRK2* KO background rescued the elevated basal ISG expression (Fig. 3A and S3A). With lowered resting type I IFN levels, cGAS/*LRRK2* double KO macrophages were able to respond normally to type I IFN-inducing innate immune stimuli like LPS and cGAMP, which bypass cGAS (Diner et al., 2013), but not ISD, which requires cGAS (Fig. 3B and S3A). Together, these results indicate that the high basal type I IFN levels in *LRRK2* KO macrophages are due to engagement of the cGAS-dependent DNA sensing pathway.

**Figure 3.**
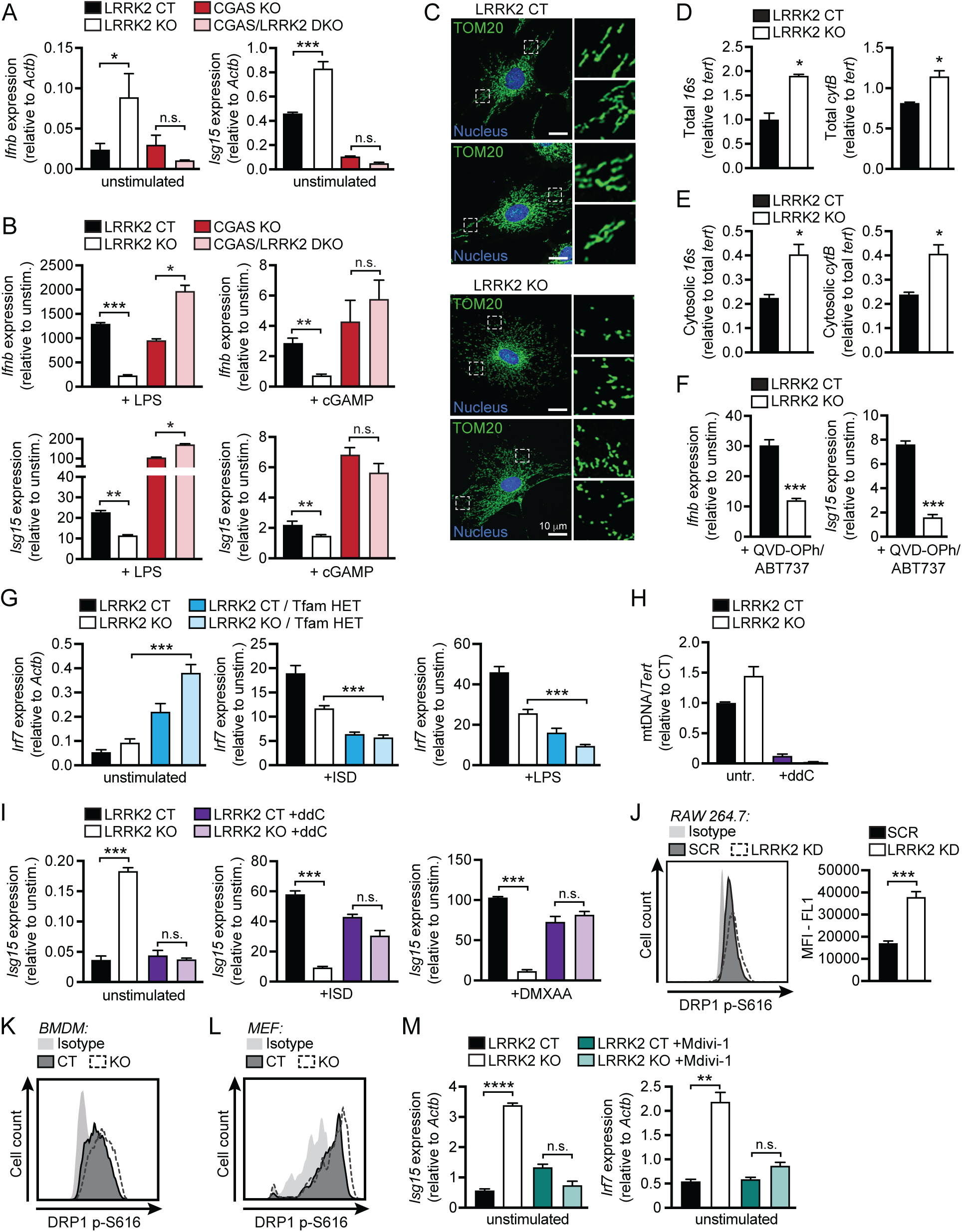
Mitochondrial DNA contributes to type I IFN dysregulation in *LRRK2* KO macrophages. **(A)** RT-qPCR of *Ifnb* and *Isg15* expression normalized to *Actb* in BMDMs from CT, *LRRK2* KO, *cGAS* KO, and *LRRK2* KO/*cGAS* double KO (DKO) mice. **(B)** As in (A) but fold-change in *Ifnb* and *Isg15* expression after transfection of 1 μg/ml ISD for 4h. **(C)** Immunofluorescence microscopy of mitochondrial networks in *LRRK2* KO vs. CT MEFs. TOM20 (green); nucleus (blue). **(D)** qPCR of total *16s* and *cytB* (mitochondrial DNA) relative to *tert* (nuclear DNA) in *LRRK2* KO vs. CT MEFs. **(E)** As in (D) but cytosolic *16s* and *cytB*. **(F)** RT-qPCR of fold-change in *Ifnb* and *Isg15* expression after treatment with 10 μM ABT737 (BCL2 inhibitor) and 10 μM QVD-OPh (caspase inhibitor) in *LRRK2* KO vs. CT BMDMs. RT-qPCR of *Irf7* expression normalized to *Actb* and fold-change in *Irf7* expression following transfection of 1 μg/ml ISD (4 h) or stimulation with 100 ng/ml LPS in BMDMs from CT, *LRRK2* KO, *Tfam* Het, and *LRRK2* KO/*TFAM* HET mice. **(H)** qPCR of ratio of *dLOOP* (mitochondrial DNA) to *Tert* (nuclear) in CT and *LRRK2* KO RAW 264.7 macrophages (normalized to CT = 1) after 4 days of 10 μM ddC treatment. **(I)** RT-qPCR of *Isg15* expression normalized to *Actb* and fold-change in *Isg15* expression following transfection of 1 μg/ml ISD (4 h) or 50 ng/ml DMXAA (2 h) in *LRRK2* KO vs. CT ddC-treated RAW 264.7 macrophages. **(J)** Histogram of counts of phospho-S616 Drp1 in *LRRK2* KD vs. SCR RAW 264.7 cells as measured by flow cytometry. **(K-L)** As in (J) but in *LRRK2* KO vs. CT (K) BMDMs and (L) MEFs. **(M)** RT-qPCR of *Isg15* and *Irf7* expression normalized to *Actb* in LRRK2 KO vs. CT BMDMs with or without 50 μM Mdivi-1 (12 h). Data represented as means +/- S.E.M. *p<0.05, **p<0.01, ***p<0.005. See also Figure S3.

### Cytosolic sensing of mtDNA contributes to basal type I IFN expression in *LRRK2* KO macrophages

We next sought to identify the source of the cGAS-activating signal. Mitochondrial DNA (mtDNA) has been shown to be a potent activator of type I IFN downstream of cGAS (Yang et al., 2014), and LRRK2 is known to influence mitochondrial homeostasis, albeit through mechanisms that are not entirely clear. To begin implicating mtDNA in type I IFN dysregulation in *LRRK2* KO cells, we first observed the status of the mitochondrial network in *LRRK2* CT and KO primary MEFs. As had previously been described for cells overexpressing wild-type or mutant alleles of *LRRK2* (Yang et al., 2014), *LRRK2* KO MEFs had a more fragmented mitochondrial network, especially around the cell periphery (Fig. 3C). We hypothesized that this fragmentation was a sign of mitochondrial damage and could allow mitochondrial matrix components, including mtDNA, to leak into the cytosol. Therefore, we isolated the cytosolic fraction of *LRRK2* CT and KO MEFs and measured cytosolic mtDNA levels. We found that *LRRK2* KO MEFs had ∼2-fold higher total mtDNA compared to controls (Fig. 3D), and importantly, *LRRK2* KO cells had ∼2-fold more cytosolic mtDNA (Fig. 3E). To exacerbate the proposed defect, we treated *LRRK2* CT and KO BMDMs with the caspase/Bcl2 inhibitor combination, Q-VD-OPh + ABT-737, which induces mitochondrial stress and spillage of mtDNA into the cytosol. As expected, this treatment specifically induced ISG expression, and consistently, lower levels of ISGs were induced in *LRRK2* KO macrophages (Fig. 3F and S3B). We next attempted to exacerbate the *LRRK2* defect by crossing *LRRK2* KO mice with *TFAM* heterozygous mice, which are deficient in the mitochondrial transcription factor required for maintaining the mitochondrial network (Kasashima et al., 2011; West et al., 2015). Indeed, depleting *TFAM* in *LRRK2* KO BMDMs further elevated basal ISG expression and amplified their inability to induce ISG expression upon innate immune stimulation (ISD or LPS) (Fig. 3G). Together, these data suggest that spillage of mtDNA into the cytosol in *LRRK2* KO cells contributes to their defective type I IFN expression.

To further implicate mtDNA in contributing to an abnormal type I IFN response in *LRRK2* KO cells, we sought to rescue the defect by depleting mtDNA using ddC (2’,3’-dideoxycytidine), an inhibitor of mtDNA synthesis, or ethidium bromide (EtBr), an intercalating agent shown to deplete mtDNA in dividing cells (Leibowitz, 1971; Meyer and Simpson, 1969). Treating *LRRK2* KO RAW 264.7 cells with ddC or EtBr substantially reduced mtDNA copy number and resulted in similar basal expression of type I IFN and ISGs in CT and KO cells (Fig. 3H-I and S3C-E). Importantly, *LRRK2* KO macrophages with depleted mtDNA had a restored type I IFN response when stimulated with ISD or DMXAA (Fig. 3I and S3D-E). This demonstrates a critical role for mtDNA in driving both the high basal levels of type I IFN and the inability to properly induce type I IFN expression in *LRRK2* KO macrophages.

Previous studies of microglia have shown that LRRK2 contributes to mitochondrial homeostasis through interaction with the mitochondrial fission protein DRP1 (Ho et al., 2018). Thus, we hypothesized that the loss of *LRRK2* may compromise mitochondrial stability via misregulation of DRP1 activity, leading to fragmented mitochondria and spillage of mtDNA into the cytosol. We observed DRP1+ puncta via immunofluorescence microscopy at the ends of fragmented mitochondria in *LRRK2* KO MEFs, but the total levels and overall distribution did not differ between the CT and KO cells (Fig. S3F). DRP1 is positively regulated via phosphorylation at Ser616 (Taguchi et al., 2007). Therefore, to measure DRP1 activity in *LRRK2* CT and KO cells, we performed flow cytometry with an antibody specific for phospho-S616 DRP1. We found increased phospho-S616 in resting *LRRK2* KD RAW 264.7 cells and *LRRK2* KO BMDMs and MEFs (Fig. 3J-L). This difference was exacerbated by the addition of H_2_O_2_, which induces DRP1-dependent mitochondrial fission, and eliminated with the addition of Mdivi-1, a specific inhibitor of DRP1 (Fig. S3G). Next, to test if DRP1 activity influences ISG expression in *LRRK2* KO cells, we chemically inhibited DRP1 with Mdivi-1 and measured basal gene expression. In *LRRK2* KO BMDMs and *LRRK2* KD RAW 264.7 macrophages, DRP1 inhibition returned ISG expression to control levels (Fig. 3M and S3H). Furthermore, DRP1 inhibition also restored the cytosolic mtDNA levels in *LRRK2* KO cells to those of CT (Fig. S3I-J). Together, these data indicate that dysregulated ISG expression in *LRRK2* KO cells is caused by leakage of mtDNA into the cytosol, which is downstream of excessive DRP1-induced mitochondrial fission.

### *LRRK2* KO macrophages are susceptible to mitochondrial stress and have altered cellular metabolism

Given that cytosolic mtDNA contributes to type I IFN defects in *LRRK2* KO macrophages, we predicted that mitochondria in *LRRK2* KO cells may be more damaged or more prone to damage. To better understand the health of the mitochondrial network in *LRRK2* KO vs. CT macrophages, we first used the carbocyanine dye JC-1, which accumulates in mitochondria to form red fluorescent aggregates. Upon loss of mitochondrial membrane potential JC-1, diffuses into the cytosol where it emits green fluorescence as a monomer. Thus, a decrease in red fluorescence (aggregates) and increase in green fluorescence (monomers) signifies mitochondrial depolarization, making JC-1 dye a highly sensitive probe for mitochondrial membrane potential. Flow cytometry analysis of resting *LRRK2* CT and KO cells revealed lower levels of JC-1 dye aggregation (i.e., lower mitochondrial membrane potential) in *LRRK2* KO BMDMs (Fig. 4A-B), *LRRK2* KD RAW 264.7 macrophages (Fig. S4A), and *LRRK2* KO MEFs (Fig. S4B). In addition, *LRRK2* KO cells were more sensitive to the mitochondrial damaging and depolarizing agents, rotenone and ATP, which is consistent with *LRRK2* KO cells harboring a baseline mitochondrial defect (Fig. 4C and S4A-B). Interestingly, the mitochondrial membrane potential of *LRRK2* KO macrophages is normalized after treatment with Mdivi-1 to inhibit DRP1, suggesting that misregulation of DRP1 is upstream of defects in mitochondrial membrane potential (Fig. 4D).

**Figure 4.**
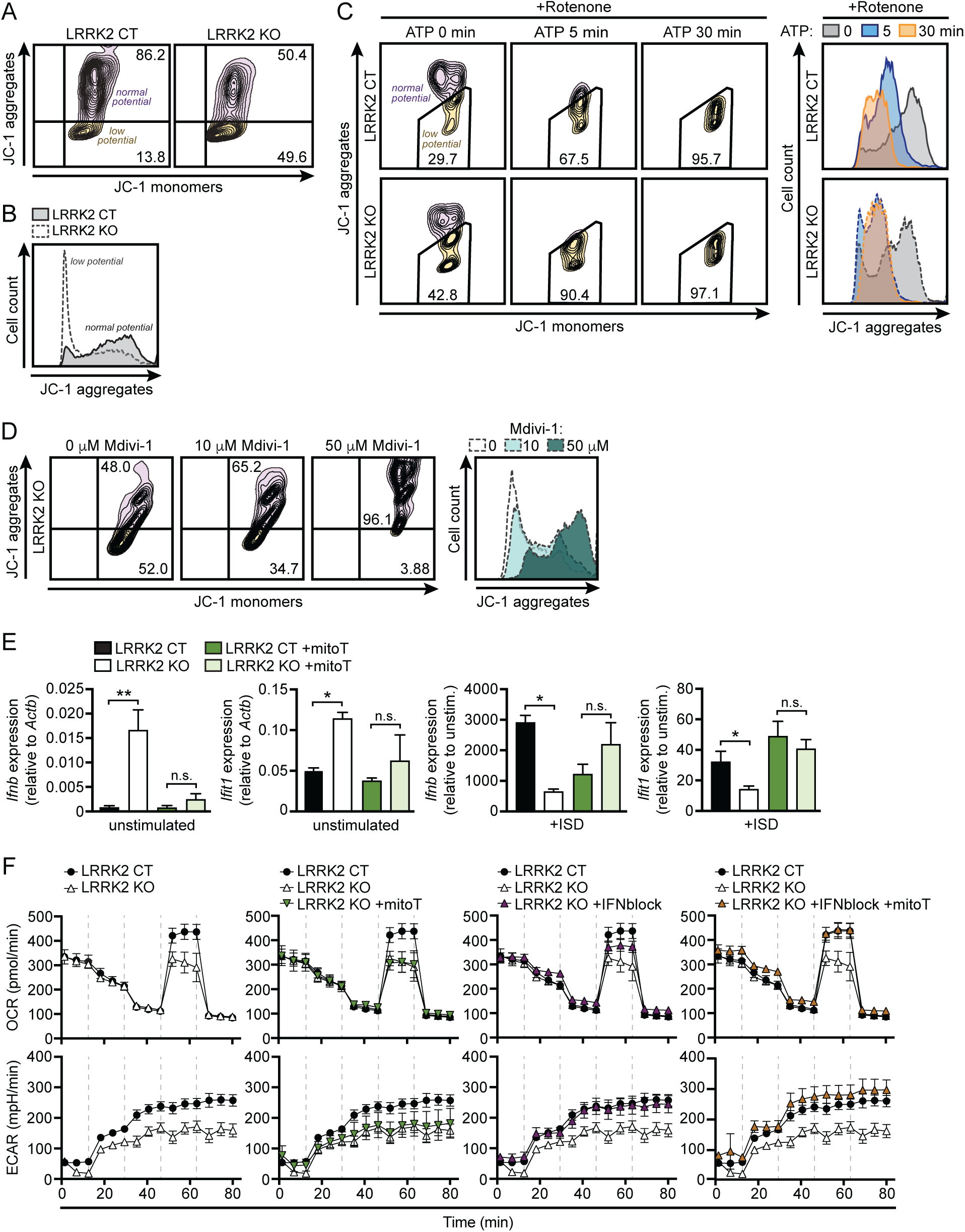
*LRRK2* KO macrophages have increased mitochondrial stress and altered metabolism. **(A)** Flow cytometry of mitochondrial membrane potential as measured by JC-1 dye aggregates (610/20) (normal membrane potential) vs. monomers (520/50) (low membrane potential) in *LRRK2* KO vs. CT BMDMs. **(B)** Histogram of cell counts and JC-1 aggregates measured by flow cytometry in *LRRK2* KO vs. CT BMDMs**. (C)** Flow cytometry of JC-1 aggregates vs. monomers and histogram of cell counts and JC-1 aggregates in *LRRK2* KO vs. CT BMDMs after treatment with 2.5 μM rotenone (3 h) followed by 5 μM ATP for indicated times. **(D)** Flow cytometry of JC-1 aggregates vs. monomers *LRRK2* KO BMDMs treated for 12 h with indicated concentration of Mdivi-1. **(E)** RT-qPCR of *Ifnb* and *Ifit1* expression normalized to *Actb* and fold-change in *Ifnb* and *Ifit1* expression following transfection of 1 μg/ml ISD (4 h) in *LRRK2* KO vs. CT BMDMs with or without 200μM mitoTEMPO (mitoT) for 16 h. **(F)** Seahorse metabolic analysis of oxygen consumption rate (OCR) and extracellular acidification rate (ECAR) in *LRRK2* KO vs. CT BMDMs untreated or treated with 200 μM mitoT, IFN-β blocking antibody, or both overnight. Data represented as means +/- S.E.M. *p<0.05, **p<0.01, ***p<0.005. See also Figure S4.

Previous reports have indicated that LRRK2 dysfunction alters reactive oxygen species (ROS) (Pereira et al., 2014; Russo et al., 2019). To test whether ROS could contribute to the defective type I IFN signature in *LRRK2* KO cells, we treated control and *LRRK2* KO BMDMs with mitoTEMPO (mitoT), a mitochondrially-targeted scavenger of superoxide (Liang et al., 2010). *LRRK2* KO cells treated with mitoT had similar ISG levels to controls both at rest and after ISD stimulation (Fig. 4E). We also hypothesized that mitochondrial defects may render *LRRK2* KO macrophages incapable of meeting metabolic demands in response to carbon sources. To test this, we altered the concentration of sodium pyruvate, an intermediate metabolite of glycolysis and the TCA cycle, in the media of *LRRK2* CT and KO BMDMs. Indeed, we observed that elevated concentrations of sodium pyruvate exacerbated high basal levels of type I IFN and decreased induction of ISGs in *LRRK2* KO macrophages in a dose-dependent manner (Fig. S4C).

Next, we further investigated the nature of the mitochondrial defect in *LRRK2* KO macrophages using the Agilent Seahorse Metabolic Analyzer. In this assay, oxidative phosphorylation (OXPHOS) and glycolysis are assayed by oxygen consumption rate (OCR) and extracellular acidification rate (ECAR), respectively. We found that OCR in *LRRK2* KO BMDMs was defective both in terms of maximal and reserve capacity (Fig. 4F, upper graph), indicating reduced mitochondrial metabolism. We also found defects in non-glycolytic acidification and in maximal and reserve capacity as measured by ECAR, indicating reduced glycolysis (Fig. 4F, lower graph). This result was surprising as cells typically switch from OXPHOS to glycolysis when activated (Kelly and O’Neill, 2015). Remarkably, co-treatment of *LRRK2* KO BMDMs with mitoT and IFN-β neutralizing antibody completely restored OCAR and ECAR. This rescue was greater than treatment of either IFN-β blockade or mitoT alone (Fig. 4F). Conversely, the addition of sodium pyruvate exacerbated these metabolic defects in *LRRK2* KO cells (Fig. S4D). Collectively, these data demonstrate that loss of *LRRK2* in macrophages has a profound impact on the mitochondria, rendering them less capable of effectively processing high-energy electrons produced by the TCA cycle.

### Reduced antioxidants and purine biosynthesis metabolites contribute to mitochondrial damage and type I IFN expression in LRRK2 KO macrophages

To better understand possible mechanisms driving or resulting from damaged mitochondria in *LRRK2* KO macrophages, we performed an unbiased query of metabolites using LC/MS/MS (Table S2) (Zhou et al., 2012). In *LRRK2* KO BMDMs, we found lower levels of inosine monophosphate (IMP) and hypoxanthine, two intermediates in the purine biosynthesis pathway, which we validated using pure molecular weight standards (Fig. 5A-C and S5A-B). Interestingly, purine metabolism is tightly associated with generation of antioxidant compounds, and several metabolites in this pathway are well-characterized biomarkers of PD (Chen et al., 2012). Consistent with lower levels of antioxidants, we detected increased oxidized glutathione and glutamate metabolism compounds in *LRRK2* KO macrophages (Table S2). Additionally, consistent with defects in purine metabolism, we observed significantly fewer puncta containing formylglycinamidine ribonucleotide synthase (FGAMS, also known as PFAS), a core purinosome component, per *LRRK2* KO cell compared to controls (Fig. 5D-E).

**Figure 5.**
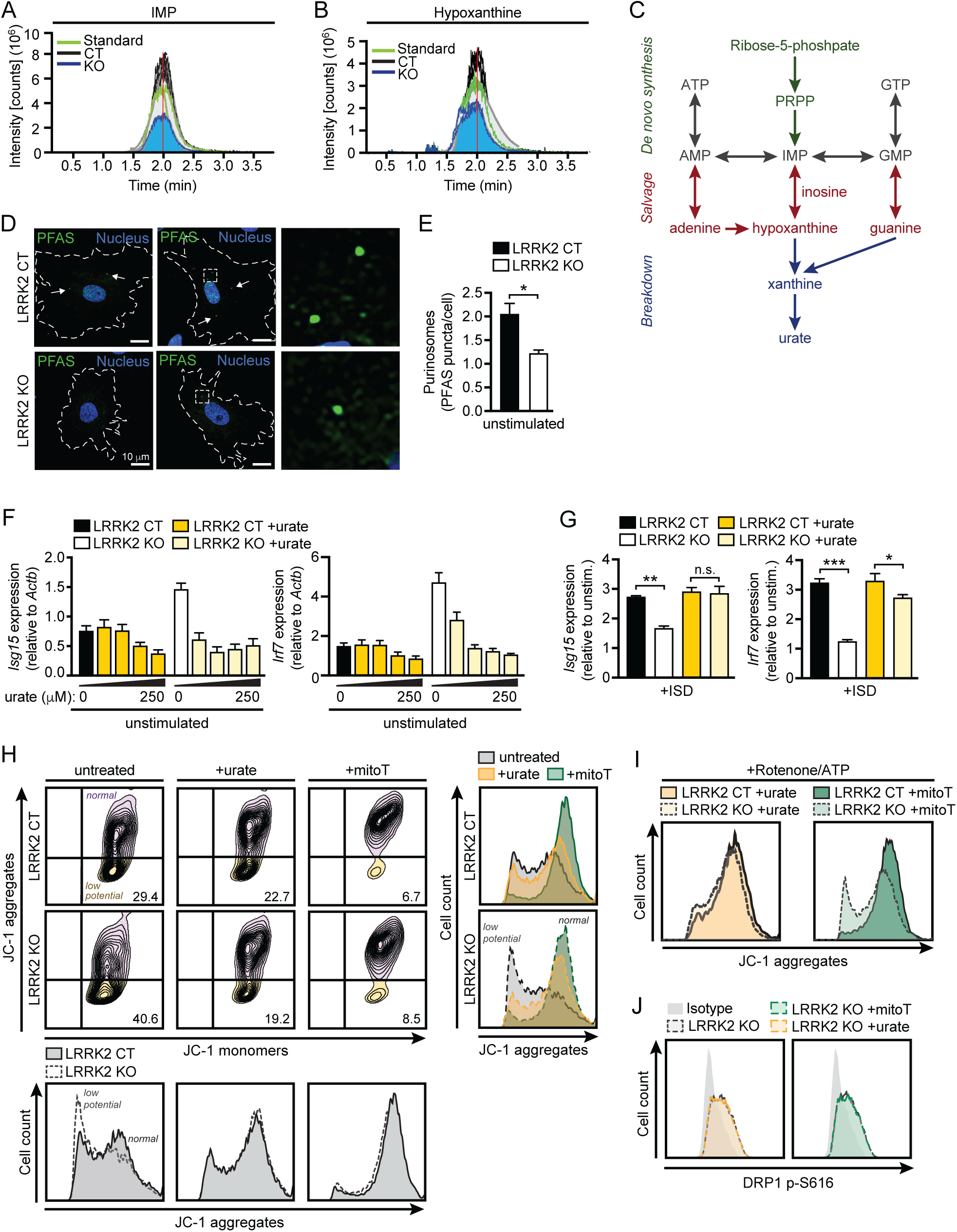
Reduced antioxidant pools in *LRRK2* KO macrophages results in mitochondrial stress. **(A)** Chromatogram of targeted metabolomic analysis of *LRRK2* KO vs. CT BMDMs with pure molecular weight standard for IMP. **(B)** As in (A) but for hypoxanthine. **(C)** Diagram of key metabolites produced during the major steps of purine metabolism. *De novo* synthesis (green), salvage (red), breakdown (blue). **(D)** Representative immunofluorescence image of purinosome formation in *LRRK2* KO vs. CT MEFs as visualized by PFAS puncta (green). Nuclei (blue). **(E)** Quantification of PFAS puncta per cell as imaged in (D). **(F)** RT-qPCR of *Isg15* and *Irf7* expression normalized to *Actb* in *LRRK2* KO vs. CT BMDMs treated with increasing concentrations of urate, (0, 10, 50, 100, and 250 μM) for 24 h. **(G)** RT-qPCR of fold-change in *Isg15* and *Irf7* expression following transfection of 1 μg/ml ISD (4 h) in *LRRK2* KO vs. CT BMDMs treated with 100 μM urate for 24 h. **(H)** Flow cytometry of JC-1 aggregates vs. monomers and histograms of cell counts and JC-1 aggregates in *LRRK2* KO vs. CT BMDMs after treatment with 100μM urate or 200μM mitoTEMPO overnight. **(I)** Histograms as in (H) but in the presence of 2.5 μM rotenone (3 h) and 2.5 μM ATP (15 min). **(J)** Histogram of counts of phospho-S616 Drp1 as measured by flow cytometry in *LRRK2* KO vs. CT BMDMs after treatment with 100μM urate or 200μM mitoTEMPO overnight. Data represented as means +/- S.E.M. *p<0.05, **p<0.01, ***p<0.005. See also Figure S5 and Table S2.

Because depleted antioxidant pools and concomitant accumulation of ROS can lead to mitochondrial damage, we hypothesized this might contribute to the mitochondrial and type I IFN defects we observe in *LRRK2* KO macrophages. To test this, we first supplemented cells with antioxidants directly in order to rescue the type I IFN defect in *LRRK2* KO macrophages. Addition of urate reduced basal ISG expression in *LRRK2* KO BMDMs (Fig 5F) and restored the ability of *LRRK2* KO BMDMs to induce ISG expression upon stimulation (Fig 5G). In addition, treatment with urate or mitoT restored the resting mitochondrial membrane potential of *LRRK2* BMDMs (Fig. 5H-I). Neither urate nor mitoT altered DRP1 activation in LRRK2 KO BMDMs, suggesting the antioxidant defects are either downstream or independent of LRRK2-dependent DRP1 misregulation (Fig. 5J). Collectively, these results suggest that the depletion of antioxidant pools in *LRRK2* KO macrophages due to defective purine metabolism contributes to their mitochondrial dysfunction and aberrant type I IFN expression.

### *LRRK2* KO mice control Mtb infection similarly to CT but have altered infection-induced neuroinflammation

Previous reports have linked SNPs in *LRRK2* with susceptibility to mycobacterial infection in humans, and our studies indicate that LRRK2 plays a key role in homeostasis of macrophages, the first line of defense and replicative niche of Mtb. Therefore, we sought to understand how LRRK2 deficiency influences innate immune responses *in vivo* during Mtb infection. We infected *LRRK2* CT and KO mice with ∼150 CFUs via aerosol chamber delivery. At 7, 21, 63, and 126 days post-infection, we observed no significant differences in bacterial burdens in the lungs or spleens of infected mice (Fig. S6A). We also measured serum cytokines and tissue cytokine expression and found no major differences (Fig. S6B-C). However, upon inspection of lung tissues via H&E staining, we observed significantly more neutrophils (polymorphonuclear leukocytes, PMNs) in the lungs of *LRRK2* KO mice 21 days post-infection (Fig. 6A-B). Furthermore, the percentage of neutrophils that were undergoing cell death (degenerate PMNs) was higher in the *LRRK2* KO mice than in the controls (Fig. 6B). The *LRRK2* KO mice also had more granulomatous nodules, indicating more macrophages had infiltrated the infected lungs (Fig. S6D-E). Together, these results indicate that while *LRRK2* CT and KO mice did not display different bacterial burdens, the *LRRK2* KO mice had disproportionate innate immune response early during Mtb infection.

**Figure 6.**
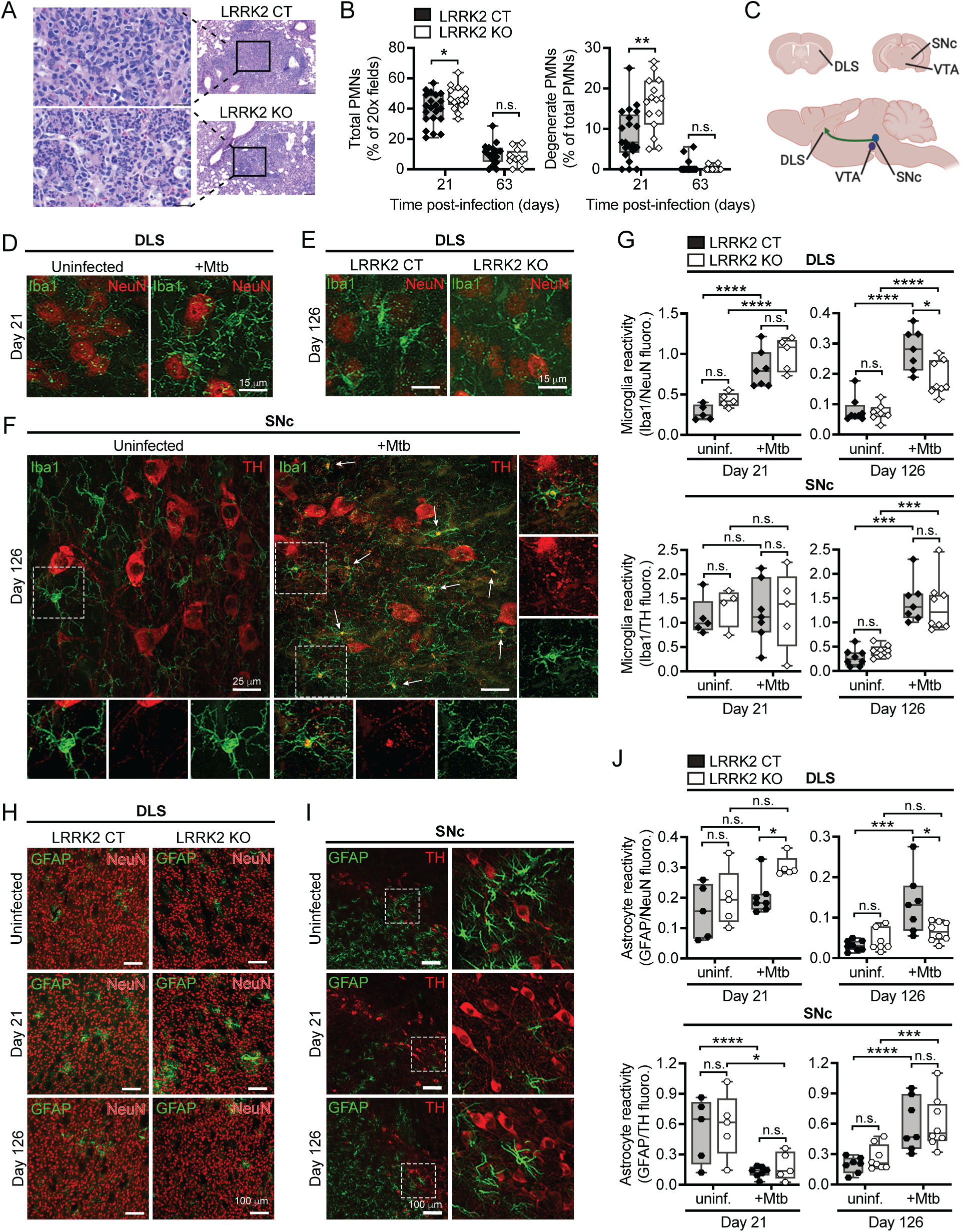
*LRRK2* KO mice exhibit increased lung pathology and activation of glial cells in the brain during Mtb infection. **(A)** Representative histology images of inflammatory nodules in the lungs of *LRRK2* KO and CT mice 21 days post-Mtb. Hematoxylin and eosin (H&E) stain. **(B)** Semi-quantification of neutrophils in the lungs of *LRRK2* KO and CT mice infected with Mtb for 21 or 63 days. Total neutrophil scores were determined by the percentage of 20x magnification fields of containing neutrophils. Degenerate neutrophil scores determined by the percentage of PMN+ fields containing degenerate neutrophils. **(C)** Schematic representation of brain areas of interest (DLS, SNc, and VTA) in the nigrostriatal dopaminergic pathway in the mouse brain. Created using BioRender. **(D-E)** Fluorescence images of reactive microglia in the DLS in *LRRK2* KO vs. CT mice uninfected or infected with Mtb for 21 or 126 days. Iba1 (green); NeuN (red). **(F)** Fluorescence images of reactive microglia in the SNc in *LRRK2* CT mice uninfected or infected with Mtb for 126 days. Iba1 (green); TH (red). Arrows highlight instances of colocalization of Iba1 and TH. **(G)** Quantification of microglial reactivity in the DLS and SNc as measured by Iba1 fluorescence relative to NeuN or TH fluorescence in *LRRK2* KO and CT mice infected with Mtb for 21 or 126 days compared to uninfected age-matched controls. **(H)** Fluorescence images of reactive astrocytes in the DLS in *LRRK2* KO vs. CT mice uninfected or infected with Mtb for 21 or 126 days. GFAP (green); NeuN (red). **(I)** As in (H) but in the SNc. GFAP (green); TH (red). **(J)** Quantification of astrocyte reactivity in the DLS and SNc as measured by GFAP fluorescence relative to NeuN or TH fluorescence in *LRRK2* KO and CT mice infected with Mtb for 21 or 126 days compared to uninfected age-matched controls. Data represented as means +/- S.E.M. *p<0.05, **p<0.01, ***p<0.005. See also Figure S6.

### Mtb infection increases microglia reactivity in PD-relevant brain regions

Several lines of evidence point to a connection between persistent infections and neurodegenerative disease, and several links between *LRRK2* and neuroinflammation have been previously reported (De Chiara et al., 2012; Schildt et al., 2019). Therefore, we set out to investigate markers of neuroinflammation in the brains of Mtb infected mice. We first focused on microglia since these are the cells of the central nervous system (CNS) that play an important role in neuroimmune surveillance, similar to macrophages in the periphery (Ousman and Kubes, 2012). To assess the extent to which Mtb infection alters microglia reactivity in *LRRK2* CT and KO mice, we focused on three brain structures relevant to PD: the dorsolateral striatum (DLS), the substantia nigra pars compacta (SNc), and the ventral tegmental area (VTA). The DLS contains dopaminergic (DA) terminals while the SNc and VTA contain DA cell bodies (Fig. 6C), and loss of these neurons is one hallmark of PD. To measure microglia reactivity, we measured endogenous Iba1 fluorescence intensity in microglia and normalized these values to neuronal nuclear protein (NeuN) fluorescence (a marker for mature neurons) in the DLS or tyrosine hydroxylase (TH) fluorescence (a marker for dopaminergic neurons) in the SNc and VTA (Fig. 6D-G and S6G-H).

We first examined the effect of systemic Mtb infection on *LRRK2* CT mice. Importantly, at all time points examined, no Mtb bacilli were detected in the brains as measured by acid-fast staining (Fig. S6F). Infected *LRRK2* CT mice showed significant increases in microglia reactivity in the DLS at 21 and 126 days post-infection compared to age-matched uninfected controls (Fig. 6D and G). In contrast, microglia reactivity in the SNc and VTA remained unaltered in *LRRK2* CT mice at 21 days post-infection but significantly increased at 126 days post-infection (Fig. 6F-G and S6H). We next examined how LRRK2 deficiency affected microglia reactivity during Mtb infection. Mtb-infected *LRRK2* KO mice showed a similar pattern of microglia reactivity in the DLS compared to *LRRK2* CT mice where microglia reactivity increased at 21 and 126 days post-infection compared to age-matched uninfected controls (Fig. 6G). Likewise, microglia reactivity in the SNc and VTA was unchanged at 21 days post-infection but increased 126 days post-infection (Fig. 6G and S6H). Because of the prolonged nature of Mtb infections, the mice in our experiments were essentially aged for an additional 126 days during the course of our experiments. Aging alone can alter the expression of Iba1 in microglia, so we examined the effect of age on baseline expression of Iba1 in *LRRK2* CT and KO mice. Both groups showed a significant age-dependent reduction in microglia reactivity in the DLS, SNc and VTA when comparing 3.5 month old to 7 month old uninfected *LRRK2* CT and KO mice (Fig. S6G). Therefore, the infection-induced increases in microglia reactivity is not due to age-related changes in Iba1 expression.

Intriguingly, upon close inspection, we observed accumulation of TH+ cellular debris and a high degree of co-localization between Iba1 and TH staining in the SNc of Mtb-infected mice 126 days post-infection (Fig. 6F, indicated by arrows). We speculate that these events correspond to phagocytosis of damaged neurons or neuronal debris by activated microglia, which appears to selectively occur in the SNc of Mtb-infected mice. Together, these data show for the first time that microglia in the the DLS, SNc, and VTA, the three PD-relevant regions of the brains, become reactive following chronic infection with Mtb. Furthermore, midbrain microglia become reactive at later time points post-Mtb infection than microglia in the DLS with no additional effect of *LRRK2* KO on the pattern or extent of reactivity. When comparing LRRK2 CT and KO mice, we did not observe any baseline differences in the reactivity profile of microglia across all brain regions or time points post-infection. The sole exception was a small but significant 1.5-fold decrease in microglia reactivity in the DLS of *LRRK2* KO mice at 126 days-post infection (Fig. 6E and G), suggesting a potential role for LRRK2 in affecting microglia reactivity specifically in the DLS late during infection.

### Loss of *LRRK2* and Mtb infection alter astrocyte reactivity in PD-relevant brain regions

We next assessed how Mtb infection alters astrocyte reactivity in *LRRK2* CT and KO mice in the DLS, SNc and VTA. Although once considered supporting cells in the CNS, emerging evidence suggests that astrocytes play vital roles in modulating neural circuit activity during physiological and pathological states (Khakh and Sofroniew, 2015). Since glial fibrillary acidic protein (GFAP) expression levels are known to increase in reactive astrocytes, we measured GFAP fluorescence and normalized values to NeuN fluorescence in the DLS and TH fluorescence in SNc and VTA (Fig. 6H-J and S6G and I).

GFAP levels in *LRRK2* CT astrocytes in the DLS were similar to uninfected mice at 21 days post-infection but significantly increased 126 days post-infection (Fig. 6H and J), suggesting chronic TB infection increases astrocyte reactivity in the DLS. In contrast to the DLS, the SNc showed a more complex profile of astrocyte reactivity. When compared to age-matched uninfected controls, SNc astrocytes showed 4-fold lower GFAP fluorescence at day 21 post-infection, but 3-fold higher GFAP fluorescence 126 days post-infection (Fig. 6I-J). The astrocyte reactivity profile within the VTA of *LRRK2* CT mice was similar to the SNc, but differences at both time points were not significant (Fig. S6I).

Having found large differences in astrocyte reactivity with systemic Mtb infection in *LRRK2* CT mice, we next assessed the reactivity of astrocytes in Mtb-infected *LRRK2* KO mice. Mtb-infected *LRRK2* KO mice had no significant changes in GFAP in the DLS 21 or 126 days post-infection compared to uninfected *LRRK2* KO mice (Fig. 6H and J). This is in contrast to the increase in GFAP fluorescence we observed for *LRRK2* CT mice at 126 days post-infection. In the SNc and VTA, however, we observed that *LRRK2* KO mice followed a similar pattern of astrocyte reactivity as *LRRK2* CT mice; astrocytes in the SNc had significantly lower GFAP at day 21 post-infection but significantly higher GFAP by day 126 post-infection (Fig. 6I-J), and GFAP fluorescence in the VTA was similar at both time points when compared to uninfected mice (Fig. S6I). Together, these data suggest that astrocytes become reactive in the DLS and SNc at later stages of Mtb infection. This pathology mirrors the progression of neurodegeneration in idiopathic PD (Stephens et al., 2005; Villalba et al., 2009; Zaja-Milatovic et al., 2005). Importantly, the dynamic reactivity of DLS astrocytes following Mtb infection is dependent on *LRRK2* expression, while SNc astrocytes do not depend on *LRRK2* for induction of reactivity following Mtb infection.

We also found that age affects astrocyte reactivity in the DLS and SNc (Fig. 6SG). Astrocyte reactivity in uninfected mice showed a significant age-dependent reduction in the DLS and SNc such that 7 month old mice showed significantly lower GFAP fluorescence than 3.5 month old mice (Fig. S6G). This age-dependent reduction was maintained in the DLS of Mtb-infected mice, but was lost in the SNc and VTA (Fig. 6J and S6I). Together these data suggest that chronic TB infection dramatically alters the reactivity profile of astrocytes in the DLS, SNc and VTA and astrocyte reactivity following Mtb infection is dependent on the expression of *LRRK2*.

### Loss of *LRRK2* impacts the ability of astrocytes to respond to stimuli *ex vivo*

Because we observed dynamic changes in astrocyte reactivity and microglial activation during the course of Mtb infection *in vivo*, we wanted to better understand the response of these cells *ex vivo*, especially in terms of the mitochondrial and type I IFN phenotypes we uncovered in *LRRK2* KO peripheral macrophages. To this end, we differentiated primary cell cultures enriched in astrocytes and microglia from the brains of neonatal *LRRK2* CT and KO mice. Astrocyte cultures were positive for *Gfap* mRNA whereas microglial cultures expressed Iba1 (Fig. 7A), which confirmed successful enrichment. *LRRK2* KO astrocyte cultures had a modest increase in *Gfap* and *Ccl5* mRNA compared to *LRRK2* CT cultures (Fig. 7B), indicating an increased reactivity at rest. Remarkably, when stimulated with IFN-β, *LRRK2* KO astrocytes selectively failed to upregulate these same markers to the extent of CT astrocytes (Fig. 7B). As we observed in *LRRK2* KO BMDMs, *LRRK2* KO astrocytes also had elevated ISG expression at rest and failed to robustly upregulate these genes upon stimulation (Fig. 7C). In addition, they were similarly more sensitive to mitochondrial depolarizing agents compared to *LRRK2* CT astrocytes (Fig 7D). Together, these results strongly suggest that *LRRK2* KO astrocytes are defective in many of the same respects we observed for peripheral macrophages.

**Figure 7.**
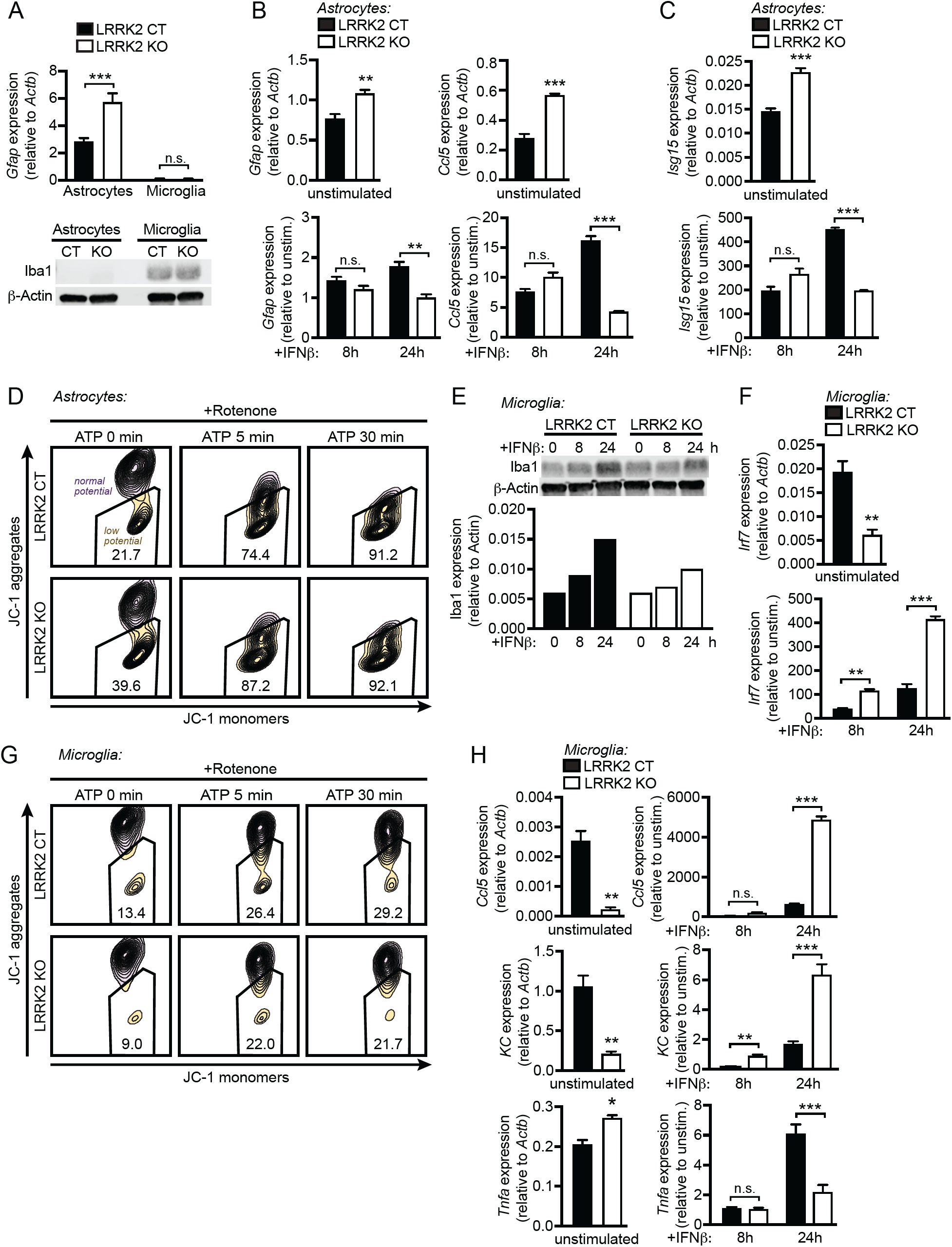
Loss of LRRK2 alters glial cell activation and reactivity *ex vivo.* **(A)** RT-qPCR of *Gfap* expression and western blot of Iba1 expression in *LRRK2* KO vs. CT astrocyte- and microglia-enriched primary cell cultures. **(B)** RT-qPCR of *Gfap* and *Ccl5* expression normalized to *Actb* and fold-change in *Gfap* and *Ccl5* expression following stimulation with 800 IU IFN-β for indicated times in *LRRK2* KO vs. CT astrocytes. **(C)** As in (B) but for *Isg15* expression. **(D)** Flow cytometry of JC-1 aggregates vs. monomers in *LRRK2* KO vs. CT astrocytes with 2.5 μM rotenone (3 h) and 2.5 μM ATP for indicated times. **(E)** Western blot of Iba1 protein levels in *LRRK2* KO vs. CT microglia treated with 800IU IFN-β. Bar graph shows quantification of protein levels relative to Actin. **(F)** RT-qPCR of *Irf7* expression normalized to *Actb* and fold-change in *Irf7* following stimulation with 800 IU IFN-β for indicated times in *LRRK2* KO vs. CT microglia. **(G)** As in (D) but for microglia. **(H)** As in (F) but for *Ccl5, KC*, and *Tnfa* expression. Data represented as means +/- S.E.M. *p<0.05, **p<0.01, ***p<0.005.

We next examined *LRRK2* KO microglia cultures *ex vivo. LRRK2* KO microglia exhibited a modest reduction in Iba1 expression upon stimulation with IFN-β compared to *LRRK2* CT cells (Fig. 7E), which is consistent with *in vivo* results in the SNc at 126 days post-Mtb infection (Fig. 6G). Surprisingly, *LRRK2* KO microglia had a reduced type I IFN signature at rest and dramatically upregulated ISGs upon stimulation (Fig. 7F). Consistent with this phenotype, *LRRK2* KO microglia were not sensitive to mitochondrial depolarizing stressors (Fig. 7G). *LRRK2* KO microglia also had significantly upregulated proinflammatory chemokines *Ccl5* and *KC* but failed to upregulate *Tnfa* to the level of *LRRK2* CT microglia (Fig. 7H), which indicates *LRRK2* KO microglia may have an alternative proinflammatory polarization compared to *LRRK2* CT cells. Together, these data suggest that *LRRK2* KO microglia and astrocytes have an altered ability to sense and respond to innate immune cues and this begins to elucidate how peripheral infection, coupled with genetic defects, may precipitate neuroinflammation.

## DISCUSSION

Despite being repeatedly associated with susceptibility to mycobacterial infection and other inflammatory disorders, very little is known about how LRRK2, a massive, multifunctional protein, functions outside of the central nervous system. Here, we provide evidence that loss of *LRRK2* influences the ability of immune cells, both in the periphery and in PD relevant regions of the brain, to respond to and express inflammatory molecules. During Mtb infection, in peripheral organs like the lungs, these defects manifest at the level of local neutrophil cell death and macrophage infiltration without significantly impacting bacterial replication. In the brains of Mtb-infected mice, loss of *LRRK2* sensitizes glial cells like astrocytes and microglia, inducing a hyper-reactive phenotype even when these cells are exposed to the same circulating cytokine milieu as control mice. Together, these results argue strongly for the “multiple-hit hypothesis” of neurodegenerative disease, whereby genetic susceptibility (e.g. loss of *LRRK2*) coupled with environmental stressors (e.g. Mtb infection (Shen et al., 2016), mitochondrial stress (Tanner et al., 2011), exhaustive exercise (Sliter et al., 2018)) can trigger neuroinflammation and potentially cause downstream damage to neurons (Balin and Appelt, 2001; Patrick et al., 2019).

We propose that dysregulation of type I IFN expression in *LRRK2* KO macrophages is the consequence of two distinct cellular defects conferred by loss of LRRK2. First, in the absence of LRRK2, decreased levels of purine metabolites and urate contribute to oxidative stress, leading to damage of the mitochondrial network. A recent human kinome screen identified LRRK2 as a kinase involved in dynamics of the purinosome, a cellular body composed of purine biosynthetic enzymes that assembles at or on the mitochondrial network (French et al., 2016). Specifically, shRNA knockdown of *LRRK2* in HeLa cells inhibited purinosome assembly and disassembly. As purinosomes are posited to form in order to protect unstable intermediates and increase metabolic flux through the *de novo* purine biosynthetic pathway (An et al., 2008; Schendel et al., 1988; Zhao et al., 2013), we propose that LRRK2-dependent defects in purinosome assembly lead to lower levels of IMP and hypoxanthine. Lower levels of these purine nucleotide intermediates in *LRRK2* KO macrophages are especially notable in the context of PD since the plasma of PD patients (both *LRRK2* mutant and idiopathic) has been shown to contain significantly less hypoxanthine and uric acid, which is the final product of the purine biosynthetic/salvage pathway. (Johansen et al., 2009). Furthermore, *LRRK2* mutation carriers with higher urate plasma levels are less likely to develop PD (Bakshi et al., 2019), and urate is currently being investigated as a potential therapeutic for PD. This highlights the importance of purine biosynthesis in the maintenance of healthy neurons.

Second, we propose that loss of *LRRK2* contributes to type I IFN dysregulation through phosphorylation of the mitochondria-associated fission protein DRP1. Previous reports have shown that LRRK2 physically interacts with DRP1 and that LRRK2 mediates mitochondrial fragmentation through DRP1 (Bakshi et al., 2019). Overexpression of both wild-type *LRRK2* and the G2019S mutant allele have been shown to cause mitochondrial fragmentation (X. Wang et al., 2012), while overexpression of the E193K allele limits mitochondrial fission by altering the LRRK2/DRP1 interaction (Carrion et al., 2018). This suggests that LRRK2 may play an important role in the formation of protein complexes, and the balance of LRRK2 expression and activity is crucial for maintenance of the mitochondrial network. Treatment of a microglia cell line (BV-2) with the LPS has previously been shown to enhance mitochondrial fission and neuroinflammation, which Ho et al. propose occurs by increasing LRRK2 and DRP1 levels (Ho et al., 2019; Perez-Carrion et al., 2018; Su and Qi, 2013; X. Wang et al., 2012). These results, coupled with our own observation that the Mtb-induced peripheral cytokine milieu can activate microglia and astrocytes, begin to paint a complex picture whereby tipping the balance of LRRK2 and DRP1 levels can trigger a pathogenic feedback loop, leading to fragmentation of mitochondria and activation of type I IFN responses. In the absence of *LRRK2*, lower antioxidant levels (via the aforementioned purinosome abnormalities) likely exacerbate this defect, leading to higher oxidative stress and further damage to the mitochondrial network.

Although we observed a striking type I IFN defect in a number of primary cells and cell lines (both higher basal levels and an inability to induce ISG expression downstream of cytosolic nucleic acid sensing or IFNAR engagement), we did not detect major differences in *in vivo* IFN-β levels in circulating serum or infected tissues at select key time points during Mtb infection. These results were surprising to us as Mtb is a potent activator of cytosolic DNA sensing (Manzanillo et al., 2012; Watson et al., 2015), and type I IFN is an important biomarker of Mtb infection associated with poor outcomes (Berry et al., 2010). We previously observed a similar apparent disconnect between type I IFN expression in vivo and ex vivo in cGAS KO mice; loss of the cytosolic DNA sensor cGAS almost completely abrogates type I IFN expression in macrophages but has only minor effects in the serum and tissues of infected mice (Watson et al., 2015). This suggests that mechanisms Consistent with our macrophage data, another recent publication investigating the role of *LRRK2* in controlling Mtb infection reported a significant decrease in IFN-α in the lungs of *LRRK2* KO infected mice 56 days post-infection (Härtlova et al., 2018).

Because our wild type and *LRRK2* KO mice were able to control Mtb replication to similar levels, the changes we observed in the brains of these mice cannot be attributed to differences in bacterial loads. They also cannot be readily attributed to differences in circulating cytokines, although it is possible that cytokine levels are significantly different at time points other than those measured (as is reported in Härtlova et al.). Therefore, it is remarkable that we observed an increase in microglial reactivity in Mtb-infected mice in the DLS, a region that is implicated in the initiation of PD (Villalba et al., 2009). Our data also suggest that reactive microglia are intimately associated with TH+ neuronal debris in the SNc of Mtb-infected mice (Fig. 6F). It is possible that the microglia are phagocytosing pieces of damaged neurons or perhaps damaging the neurons themselves. In either case, the fact that this behavior is only apparent in the brains of Mtb-infected mice strongly suggests that peripheral infection alters interactions between microglia and neurons in potentially pathologic ways. It is tempting to speculate that this increase in reactive microglia serves as a mechanism by which persistent infections can precipitate neurodegeneration (Patrick et al., 2019).

In contrast to microglial activation, Mtb infection induces DLS astrocyte reactivity in CT mice but not *LRRK2* KO mice. Combined with our *ex vivo* observations, this indicates that particularly astrocytes in the DLS depend on LRRK2 to initiate reactivity. Indeed, *LRRK2* mRNA expression in astrocytes in mice is ∼6 fold higher than in microglia (Y. Zhang et al., 2014). Interestingly, in the SNc we only observed infection-induced increases in astrocyte reactivity later during Mtb infection (126 days). Considering we found that loss of LRRK2 increases mitochondrial fragmentation, this strongly suggests that the differential reactivity of astrocytes in the DLS and SNc is due to differences in astrocytes’ mitochondria, and perhaps astrocytic mitochondria in the SNc are more resilient and/or functionally different from astrocytic mitochondria in the DLS.

The data presented here are the first to directly connect *LRRK2*’s role in maintaining mitochondrial homeostasis to its emerging role in regulating inflammation in both the brain and periphery. Given the striking phenotypes we observed in the brains of Mtb-infected *LRRK2* KO mice, it is tempting to speculate that LRRK2’s contribution to neuroinflammation and glial cell activation is a major driver of PD, thus opening the door for novel immune-targeted therapeutic interventions designed to halt or slow neurodegenerative disease progression.

## ACKNOWLEDGEMENTS

We’d like to thank Cory Klemashevich at the TAMU Integrated Metabolomics Analysis Core for his help with the metabolomics analysis. We would also like the acknowledge Monica Britton at the University of California, Davis DNA Technologies & Expression Analysis Core Library for her help with the RNA-seq analysis. We would like to acknowledge the members of the Patrick and Watson labs for their helpful discussions and feedback as well as Elizabeth Case for her help proofreading. We’d like to thank A. Phillip West and the West lab for their help with mitochondrial experiments and for providing us with *Tfam* Het and *cGAS* KO mice. We’d lastly like to thank Nevan Krogan at University of California, San Francisco for his help with the conceptual design of this manuscript.

This work was supported by funds from the Michael J. Fox Foundation for Parkinson’s Research, grant 12185 (to R.O.W.), the Nation Institutes of Health, grant 1R01AI125512-01A1 (to R.O.W.), and the Texas A&M Clinical Science and Translational Research (CSTR) Pilot Grant Program (to R.O.W., K.L.P., and R.S.). Additional funding was provided by the Parkinson’s Foundation Postdoctoral Fellowship (to C.G.W.), the NSF Graduate Research Fellowship Program (to T.E.H.), NIH training grant 5T32OD011083-10 (to K.J.V.), and the Texas A&M CVM Postdoctoral Trainee Research Training Grant (to K.J.V.).

## AUTHOR CONTRIBUTIONS

Conceptualization, K.L.P., R.O.W., C.G.W., S.L.B., and R.S.; Investigation, C.G.W., S.L.B., K.J.V., T.E.H., and R.O.W.; Methodology, C.G.W., S.L.B., K.J.V., R.O.W., K.L.P., T.E.H., and R.S.; Writing, K.L.P., C.G.W., S.L.B., R.O.W., T.E.H., and R.S.; Visualization, S.L.B., C.G.W., and R.O.W.; Funding acquisition, R.O.W., K.L.P., C.G.W., K.J.V., R.S., and T.E.H.; Supervision, R.O.W., K.L.P., and R.S..

## METHODS

### Mice

*LRRK2* KO mice (C57BL/6-Lrrk2^tm1.1Mjff^/J) stock #016121, and *IFNAR* KO mice (B6(Cg)-Ifnar1^tm1.2Ees^/J) stock #028288 were purchased from The Jackson Laboratories (Bar Harbor, ME). *Tfam* HET (Woo et al., 2012) and *cGAS* KO (B6(C)-Cgastm1d(EUCOMM)Hmgu/J) mice were provided by A. Phillip West, Texas A&M Health Science Center (Bryan, TX). All mice used in experiments were compared to age- and sex-matched controls. In order to ensure littermate controls were used in all experiments *LRRK2* KO crosses were made with (KO) *LRRK2*^−/−^ x (HET) *LRRK2*^+/−^ mice. Mice used to generate BMDMs and PEMs were between 8-12 weeks old. Mice were infected with Mtb at 10 weeks. Mice used to make glial cultures were P0.5 days old. Embryos used to make primary MEFs were 14.5 days post coitum. All animals were housed, bred, and studied at Texas A&M Health Science Center under approved Institutional Care and Use Committee guidelines.

### Mycobacterial infections

The Erdman strain was used for all Mtb infections. Low passage lab stocks were thawed for each experiment to ensure virulence was preserved. Mtb was cultured in roller bottles at 37°C in Middlebrook 7H9 broth (BD Biosciences) supplemented with 10% OADC, 0.5% glycerol, and 0.1% Tween-80 or on 7H11 plates (Hardy Diagnostics). All work with Mtb was performed under Biosafety Level 3 (BSL3) containment using procedures approved by the Texas A&M University Institutional Biosafety Committee.

To prepare the inoculum, bacteria grown to log phase (OD 0.6-0.8) were spun at low speed (500g) to remove clumps and then pelleted and washed with PBS twice. Resuspended bacteria were briefly sonicated and spun at low speed once again to further remove clumps. The bacteria were diluted in DMEM + 10% horse serum and added to cells at an MOI of 10. Cells were spun with bacteria for 10 min at 1000g to synchronize infection, washed twice with PBS, and then incubated in fresh media. RNA was harvested from infected cells using 0.5-1.0 ml Trizol reagent 4 h post-infection unless otherwise indicated.

*M. leprae* was cultivated in the footpads of nude mice and generously provided by the National Hansen’s Disease Program. Bacilli were recovered overnight at 33°C, mixed to disperse clumps and resuspended in DMEM + 10% horse serum. Cells were infected as with Mtb but with an MOI of 50.

### Mouse infections

All infections were performed using procedures approved by Texas A&M University Institutional Care and Use Committee. The Mtb inoculum was prepared as described above. Age- and sex-matched mice were infected via inhalation exposure using a Madison chamber (Glas-Col) calibrated to introduce 100-200 CFUs per mouse. For each infection, approximately 5 mice were euthanized immediately, and their lungs were homogenized and plated to verify an accurate inoculum. Infected mice were housed under BSL3 containment and monitored daily by lab members and veterinary staff.

At the indicated time points, mice were euthanized, and tissue samples were collected. Blood was collected in serum collection tubes, allowed to clot for 1-2 hr at room temperature, and spun to separate serum. Serum cytokine analysis was performed by Eve Technologies (Calgary, Alberta, Canada). Organs were divided to maximize infection readouts (CFUs: left lobe lung and ½ spleen; histology: 2 right lung lobes and ¼ spleen; RNA: 1 right lung lobe and ¼ spleen). For histological analysis organs were fixed for 24 h in either neutral buffered formalin and moved to ethanol (lung, spleen) or 4% paraformaldehyde and moved to 30% sucrose (brain). Organs were further processed as described below. For cytokine transcript analysis, organs were homogenized in Trizol Reagent, and RNA was isolated as described below. For CFU enumeration, organs were homogenized in 5 ml PBS + 0.1% Tween-80, and serial dilutions were plated on 7H11 plates. Colonies were counted after plates were incubated at 37°C for 3 weeks.

### Histopathology

Lungs and spleens were fixed with paraformaldehyde, subjected to routine processing, embedded in paraffin, and 5-μm sections were cut and stained with hematoxylin and eosin (H&E) or acid-fast stain (Diagnostic BioSystems). A boarded veterinary pathologist performed a masked evaluation of lung sections for inflammation using a scoring system: score 0, none; score 1, up to 25% of fields; score 2, 26-50% of fields; score 3, 51-75% of fields; score 4, 76-100% of fields. To quantify the percentage of lung fields occupied by inflammatory nodules, scanned images of at least 2 sections of each lung were analyzed using Fiji Image J (Schindelin et al., 2012) to determine the total cross-sectional area of inflammatory nodules per total lung cross sectional area. For acid fast staining, one brain hemisphere was fixed with paraformaldehyde for 48 hours, then transferred to a cryoprotective buffer (30% sucrose in a phosphate buffer), and frozen for coronal slicing into 40-μm sections. At least two sections per mouse were stained with an acid-fast stain (Diagnostic BioSystems) according to the manufacturer’s instructions and visualized by an Olympus BH2 light microscope.

### Tissue immunohistochemistry

At indicated time points, infected or uninfected mice were anesthetized with isoflurane and quickly decapitated. The brain was gently removed from the skull and postfixed in 4% paraformaldehyde overnight at 4°C. The tissue was cryoprotected in 30% sucrose + PBS solution for 48-72 hours. 40 µm thick coronal sections were obtained using a cryostat microtome (Leica) and preserved in 0.01% sodium azide + PBS at 4°C.

Immunohistochemistry was performed using previously published techniques (Srinivasan et al., 2015; 2016). Briefly, sections were washed three times for 10 min in 1X TBS, then blocked for 1 h in 5% normal goat serum (NGS) and 0.25% Triton-X-100 in 1X TBS at RT. Sections were incubated overnight at 4°C in primary antibodies diluted in blocking solution. The following primary antibodies were used: rabbit anti-GFAP (1:1000; Abcam ab7260), rabbit anti-Iba1 (1:250; Wako Chemical 019-19741), mouse anti-NeuN (1:500; Abcam ab104224), and chicken anti-TH (1:1000; Abcam ab76442). The following day, sections were washed 3x for 10 min each in 1X TBS and incubated with appropriate secondary antibodies in blocking solution for 2 h at RT. The following secondary antibodies were used: goat anti-rabbit (1:1000; Abcam ab150077), goat anti-mouse (1:1000; Abcam ab150120), and goat anti-chicken (1:1000; Abcam ab150176). The sections were rinsed 3x for 10 min in 1X TBS and then mounted on microscope slides in Fluoromount (Diagnostic Biosystems; K024) for imaging.

### Tissue imaging and analysis

Images were obtained using a FV 1200 Olympus inverted confocal microscope equipped with 20x, 0.85 NA oil immersion objective, 473 nm, and 561 nm laser lines to excite appropriate Alexa Fluor secondary antibodies. Images were obtained at 1x digital zoom. HV, gain, and offset was adjusted so that fluorescent signals from images were just below saturation. Laser power for 473 and 561 excitation lines were maintained between 2-3% of maximum. All images were acquired as z-stacks with a 1 µm step size and stack sizes ranged between 25-30 µm. Parameters were kept constant for all mice in an experimental group, which was defined based on infection status. Images were collected and processed with mouse genotypes blinded.

Images were processed using ImageJ. For image analysis, maximum intensity projections of z-stacks were first obtained. Projected images were thresholded such that GFAP staining in astrocytic cell bodies or Iba-1 staining in microglial cell bodies along with branches (1° and 2°) were masked and ROIs were obtained in this way. In each case, corresponding NeuN labeled or TH labeled sections were processed in a similar manner to astrocytic and microglial staining. Integrated density values were extracted from astrocytic, microglial, and corresponding neuronal components of each slice. Ratios of astrocytic or microglial integrated density to respective neuronal integrated density (NeuN/TH) were obtained. Ratios obtained in this way were averaged across each brain region and all slices for each mouse. By utilizing ratios of glial signal to neuronal staining intensity, we controlled for differences between individual sections that occur due to variations in the efficiency of antibody binding or tissue quality. Data are presented as averages for each mouse. Mean values ± s.e.m. from the averages are presented.

### Primary cell culture

Bone marrow derived macrophages (BMDMs) were differentiated from bone marrow (BM) cells isolated by washing mouse femurs with 10ml DMEM. Cells were then centrifuged for 5 min at 1000 rpm and resuspended in BMDM media (DMEM, 20% FBS (Millipore), 1mM Sodium pyruvate, 10% MCSF conditioned media). BM cells were counted and plated at 5×10^6^ in 15cm non-TC treated dishes in 30ml complete media. Cells were fed with an additional 15ml of media on day 3. Cells were harvested on day 7 with 1X PBS + EDTA.

Mouse embryonic fibroblasts (MEFs) were isolated from embryos. Briefly, embryos were dissected from yolk sacs, washed 2 times with cold 1X PBS, decapitated, and peritoneal contents removed. Headless embryos were disagreggated in cold 0.05% trypsin-EDTA (Lonza) and incubated on ice for 20 min and at 37°C for an additional 20 min. Cells were DNAse treated with 4ml dissagregation media (DMEM, 10% FBS, 100ug/ml DNAse) for 20min at 37°C. Cells were pelleted and resuspended in complete media (DMEM, 10% FBS, 1mM sodium pyruvate) and plated in 15cm dishes at embryo per dish. MEFs were allowed to expand for 2-3 days before harvest with Trypsin + 0.05% EDTA.

Mixed glial cultures were differentiated from the brains of neonatal mice as described (Lian et al., 2016). Microglial cells were differentiated using complete media (DMEM, 10% FBS, 1mM sodium pyruvate, 10% MCSF conditioned media).

Peritoneal macrophages (PEMs) were elicited by intraperitoneal injection of 1ml 3% Thioglycollate broth (BD Biosciences) for 4 days prior to harvest. For harvest, PEMs were isolated from mice by lavage (1X PBS 4°C) and resuspended in RPMI 1640 media with 20% FBS, 1mM sodium pyruvate, and 2mM L-Glutamine. Following overnight incubation at 37°C, cells were washed twice (1X PBS 37°C) to remove non-adherant cells (∼25% of population).

### Cell lines and treatments

RAW 264.7 LRRK2 KO cells (ATCC® SC-6004™) generated by the MJFF, were obtained from the ATCC and used with wild type control LRRK2 parental RAW 264.7 (ATCC® SC-6003™). To deplete mtDNA, RAW 264.7 cells were seeded at 2×10^6^ cells/well in 10 cm non-TC treated dishes and cultured for 4 days in complete media (DMEM, 10% FBS, 1mM sodium pyruvate) with 300 ng/ml ethidium bromide or 10μM ddC. Cells were split and harvested with 1X PBS + EDTA.

### Cell stimulations

BMDMs were plated in 12-well dishes at 5×10^5^ cells/well, or 6-well dishes at 1×10^6^ cells/well. MEFs were plated in 12-well dishes at 3×10^5^ cells/well. PEMs were plated in 24-well dishes at 1×10^6^ cells/well. RAW 264.7 cells were plated in 12-well dishes at 5×10^5^ cells/well. Astrocyte cultures were plated at 2.5×10^4^ cells/well in 12-well dishes. Microglia were plated at 5×10^5^ cells/well in 12-well dishes. U937 monocytes were plated at in 6-well dishes 1×10^6^ cells/well, cultured with 10ng/ml phorbol 12-myristate 13-acetate (PMA) for 48 h to induce differentiation, and then recovered in fresh media for an addition 24 h.

Cells were stimulated for 4 h with 1 μM CLO97, 100 ng/ml LPS, 10 μM ABT737/10 μM QVD-OPh, or transfected 1 μg/ml ISD, 1 μg/ml poly(I:C), 1 μg/ml cGAMP with lipofectamine or 1 μM CpG 2395 with Gene Juice. Macrophages were stimulated for 2 h with 10 μM DMXAA or 200 IU IFN-β. Glial cells were stimulated for 8 or 24 h with 800 IU IFN-β.

### mRNA sequencing

RNA was isolated using PureLink RNA mini kits (Ambion) and quantified on a Bioanalyzer 2100 (Agilent). PolyA+ PE 100 libraries were sequenced on a HiSeq 4000 at the UC Davis Genome Center DNA Technologies and Expression Analysis Core. Heatmaps were generated by performing Cluster analysis (Cluster3) followed by Java TreeView. Transcriptome analysis was performed using IPA analysis to generate GO term and disease pathway lists. Instant Clue was used to generate scatter plots and volcano plots.

### qRT-PCR

RNA was isolated using Directzol RNAeasy kits (Zymogen). cDNA was made with iScript Direct Synthesis kits (BioRad) per manufacturer’s protocol. qRT-PCR was performed in triplicate using Sybr Green Power-up (ThermoFisher). Data was analyzed on a ViiA 7 Real-Time PCR System (Applied Biosystems).

### Cytosolic DNA isolation

MEFs were plated in 10cm dishes at 3×10^6^. The next day, confluent plates were treated as indicated with inhibitors. To harvest, cells were lifted with PBS+EDTA. To determine total DNA content, 1% of the input was saved and processed by adding NaOH to 50 mM, boiling 30 min, and neutralizing with 1:10 1 M Tris pH 8.0. To isolate cytosolic DNA, the cells were pelleted and resuspended in digitonin lysis buffer (150 mM HEPES pH 7.4, 50 mM NaCl, 10 mM EDTA, 25 ug/ml digitonin). Cells were incubated for 15 at 4° on an end-over-end rotator. Cells were spun at 980g for 3 min, and the DNA from the supernatant (cytosolic fraction) was then extracted via phenol:chloroform (1:1 supernatant:phenol/chloroform). The DNA from the aqueous layer was precipitated in 0.3M sodium acetate, 10 mM magnesium chloride, 1ug/ml glycogen, and 75% ethanol. After freezing overnight at −20°C, the DNA was pelleted, washed in 70% ethanol, dried, resuspended in TE, and solubilized at 50°C for 30 min. qPCR was performed on the input (1:50 dilution) and cytosolic (1:2 dilution) samples using nuclear (*Tert*) and mitochondrial (*16s* and *cytB*) genes. The total and cytosolic mitochondrial DNA was normalized to nuclear DNA in order to control for variation in cell number.

### Western blot

Cells were washed with PBS and lysed in 1X RIPA buffer with protease and phosphatase inhibitors (Pierce). DNA was degraded using 1U/ml benzonase (EMD Millipore). Proteins were separated by SDS-PAGE and transferred to nitrocellulose membranes. Membranes were blocked for 1 h at RT in Odessy blocking buffer (Licor) and incubated overnight at 4°C with the following antibodies: IRF3 (Cell Signaling) 1:1000; pIRF3 Ser396 (Cell Signaling 4947) 1:1000; Iba1 (Wako Chemical 019-19741), 1:2000; Beta Actin (Abcam), 1:5000; and tubulin (abcam), 1:5000. Membranes were incubated with appropriate secondary antibody (Licor) for 2 h at RT prior to imaging on Odyssey Fc Dual-Mode Imaging System (Licor).

### Seahorse metabolic assays

Seahorse XF Mito Stress test kits and cartridges (Agilent) were prepared per manufacturers protocol and as previously described (Bossche et al., 2015). BMDMs were seeded at 5×10^4^ cells/well and analyzed the following day on a Seahorse XF 96well Analyzer (Agilent).

### Immunofluorescence microscopy

MEFs were seeded at 1×10^5^ cells/well on glass coverslips in 24-well dishes. Cells were fixed in 4% paraformaldehyde for 10 min at RT and then washed three times with PBS. Coverslips were incubated in primary antibody diluted in PBS + 5% non-fat milk + 0.1% Triton-X (PBS-MT) for 3 h. Cells were then washed three times in PBS and incubated in secondary antibodies and DAPI diluted in PBS-MT for 1 h. Coverslips were washed twice with PBS and twice with deionized water and mounted on glass slides using Prolong Gold Antifade Reagent (Invitrogen).

### Flow cytometry

#### JC-1 assay to assess mitochondrial membrane potential

Cells were lifted off culture plates with 1X PBS + EDTA (BMDMs, RAW 264.7 and microglia) or Accutase (Biolegend) (MEFs and astrocytes). Single cell suspensions were made in 1X PBS 4% FBS. JC-1 dye (ThermoFisher) was sonicated for 5 minutes with 30 second intervals. Cells were stained for 30 min at 37°C in 1 μM JC-1 dye and analyzed on an LSR Fortessa X20 (BD Biosciences). Aggregates were measured under Texas Red (610/20 600LP) and monomers under FITC (525/50 505LP). To assess mitochondrial membrane potential under stress, cells were treated for 3 h with 2.5 μM rotenone prior to being lifted of the culture plates. 5 μM ATP was then added for 5, 15, or 30 min. For rescue assays cells were treated overnight with mitoTEMPO (Sigma Aldrich) or urate (Sigma Aldrich).

#### Phospho-DPR1 assay

Cells were washed once in 1X PBS and fixed in 4% cold PFA for 10 min. Cells were then permeabilized with 0.3% Triton-X for 15 min followed by 30 min block in 0.1% Triton-X + 5% normal rat serum (Stem Cell Technologies). Cells were incubated in Drp1 p616 Ab overnight at 4°C in 0.1% Triton-X + 1% BSA and then in secondary antibodies (AF488 Goat anti-Rabbit). Cells were analyzed on an LSR Fortessa X20 (BD Biosciences) under FITC (525/50 505LP). For rescue and exacerbation assays, cells were treated with 100 μM H_2_O_2_ for 1 h at 37°C or for 12 h with 50 μM Mdivi-1 (Abcam).

### LCM/MS/MS

#### Sample Extraction

Samples were weighed and extracted with a methanol:chloroform:water based extraction method. Briefly 800 μL ice cold methanol:chloroform (1:1, v:v) was added to samples in a bead based lysis tube (Bertin, Rockville, MD). Samples were extracted on a Precyllys 24 (Bertin) tissue homogenizer for 30 seconds at a speed of 6000. The supernatant was collected, and samples were homogenized a second time with 800 μL ice methanol:chloroform. 600 μL ice cold water was added to the combined extract, vortexed and centrifuged to separate the phases. The upper aqueous layer was passed through a 0.2 μm nylon filter (Merck Millipore, Burlington, MA). 500 μL of the filtered aqueous phase was then passed through a 3 kDa cutoff column (Thermo Scientific) and the flow through was collected for analysis.

#### Sample Analysis

Untargeted liquid chromatography high resolution accurate mass spectrometry (LC-HRAM) analysis was performed on a Q Exactive Plus Orbitrap mass spectrometer (Thermo Scientific, Waltham, MA) coupled to a binary pump HPLC (UltiMate 3000, Thermo Scientific). For acquisition, the Sheath, Aux and Sweep gasses were set at 50, 15 and 1 respectively. The spray voltage was set to 3.5 kV (Pos) or 2.8 kV (Neg) and the S-lens RF was set to 50. The source and capillary temperatures were both maintained at 350°C. Full MS spectra were obtained at 70,000 resolution (200 m/z) with a scan range of 50-750 m/z. Full MS followed by ddMS2 scans were obtained at 35,000 resolution (MS1) and 17,500 resolution (MS2) with a 1.5 m/z isolation window and a stepped NCE (20, 40, 60). Samples were maintained at 4°C before injection. The injection volume was 10 µL. Chromatographic separation was achieved on a Synergi Fusion 4µm, 150 mm x 2 mm reverse phase column (Phenomenex, Torrance, CA) maintained at 30°C using a solvent gradient method. Solvent A was water (0.1% formic acid). Solvent B was methanol (0.1% formic acid). The gradient method used was 0-5 min (10% B to 40% B), 5-7 min (40% B to 95% B), 7-9 min (95% B), 9-9.1 min (95% B to 10% B), 9.1-13 min (10% B). The flow rate was 0.4 mL/min. Sample acquisition was performed Xcalibur (Thermo Scientific). Data analysis was performed with Compound Discoverer 2.1 (Thermo Scientific).

### Statistical analysis

All data are representative of 2 or more independent experiments with an n=3 or 4. Graphs were generated using Prism (GraphPad). Significance for assays were determined using a student’s two-tailed t test, or a one way ANOVA followed by a Bonferroni’s multiple comparisons test for more than two variables, unless otherwise noted. Error bars represent SEM. Statistical tests for brain sections were run in OriginPro 2019. For each experimental group 5-8 mice were utilized. Eight sections per mouse were used for respective antibody combinations 1) GFAP/NeuN 2) Iba-1/NeuN 3) GFAP/TH and 4) Iba-1/TH, for 2 sections per combination. Resulting in a total of 10-16 brain sections, which represent all of the mice within an experimental group. Utilizing the total number of sections as the sample size for each experimental group, we obtained a power of 1. For statistical comparison, each experimental group was tested for normal distribution. Normally distributed sets of data were compared using student’s two-tailed *t* test, while non-normally distributed data were tested using a two-tailed Mann Whitney’s test. Difference were considered statistically significant if p < 0.05.

