## Supplemental Figures for "LRRK2 regulates innate immune responses and neuroinflammation during *Mycobacterium tuberculosis* infection"

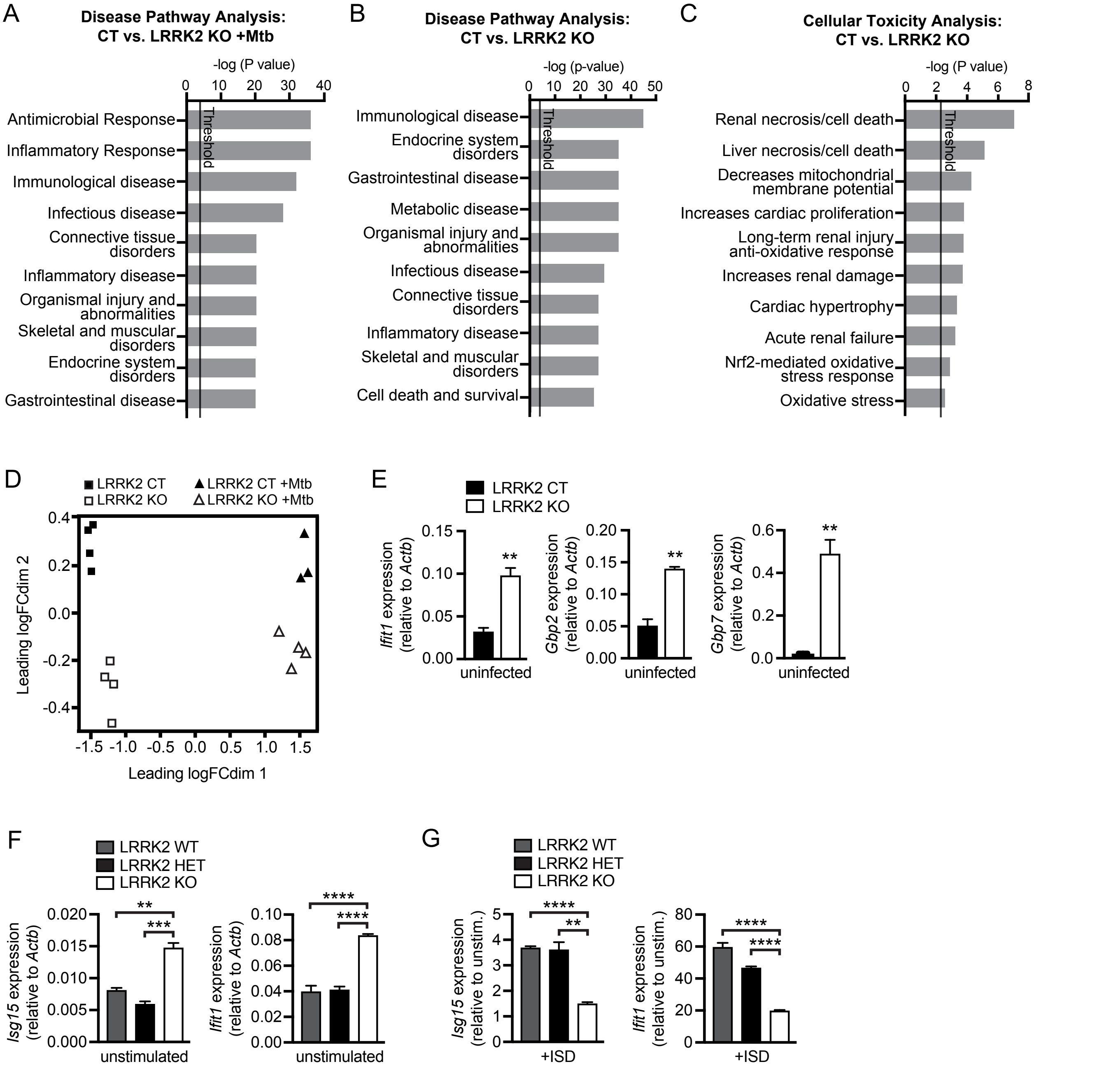


**Figure S1. (A-B)** Results of analysis performed using IPA software for top disease pathways associated with gene expression differences between (A) Mtb-infected and (B) uninfected *LRRK2* KO and CT BMDMs. **(C)** As in (B) but for pathways associated with cellular toxicity. **(D)** Principle component analysis of *LRRK2* KO vs. CT BMDMs uninfected and infected with Mtb. **(E)** RT-qPCR of type II IFN genes (*Ifit1*, *Gbp2*, and *Gbp7)* normalized to *Actb* in uninfected LRRK2 KO vs. CT BMDMs. **(F)** RT-qPCR of *Isg15* and *Ifit1* expression normalized to *Actb* in untreated wild-type (WT), *LRRK2* HET (CT), and *LRRK2* KO BMDMs. **(G)** As in (F) but fold-change in *Isg15* and *Ifit1* expression following transfection of 1 mg/ml ISD (4 h). Data represented as means +/- S.E.M. *p<0.05, **p<0.01, ***p<0.005.

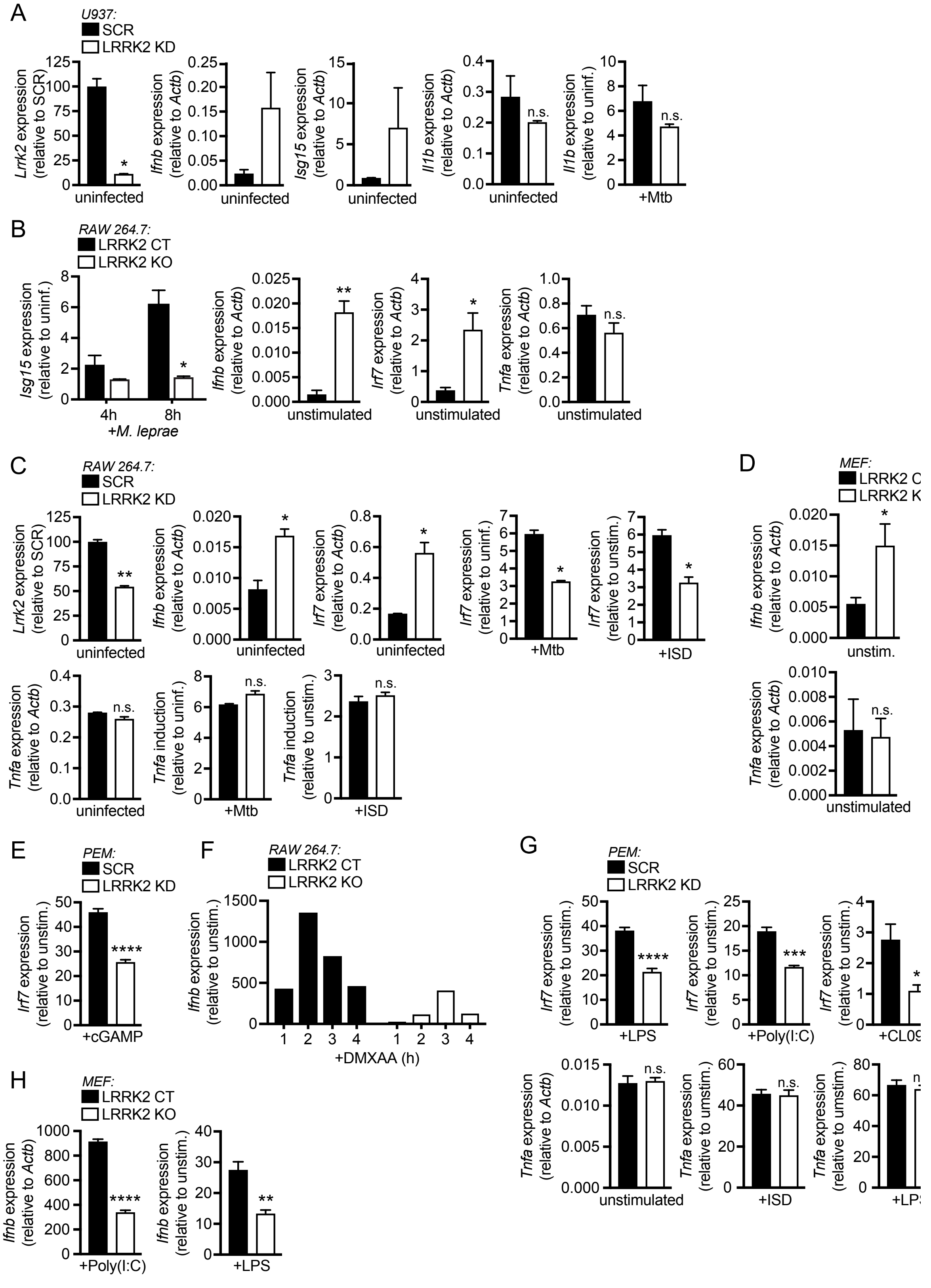


**Figure S2. (A)** RT-qPCR analysis of *LRRK2* KD U937 monocytes (from left to right): *Lrrk2* expression normalized to *Actb*; *Ifnb* and *Isg15* expression normalized to *Actb* in uninfected cells; *Il1b* expression normalized to *Actb*; fold-change in *Il1b* expression during Mtb infection (4 h). **(B)** RT-qPCR analysis of LRRK2 KO RAW 264.7 macrophages (from left to right): fold-change in *Isg15* expression during *M. leprae* infection at indicated times; *Ifnb, Irf7,* and *Tnfa* expression normalized to *Actb* in uninfected cells. **(C)** RT-qPCR analysis of *LRRK2* KD RAW 264.7 macrophages (from left to right): *LRRK2* expression normalized to *Actb*; *Ifnb* and *Irf7* expression normalized to *Actb* in uninfected cells; fold-change in *Irf7* during Mtb infection (4 h) or following transfection with 1 µg/ml ISD (4 h); *Tnf* expression normalized to *Actb* in uninfected cells; fold-change in *Tnfa* during Mtb infection (4 h) or following transfection with 1 µg/ml ISD (4 h). **(D)** RT-qPCR analysis of *LRRK2* KO MEFs (from top to bottom): *Ifnb* and *Tnfa* expression normalized to *Actb* in untreated cells. **(E)** RT-qPCR of fold-change in *Irf7* in *LRRK2* KO vs. CT PEMs transfected with 1 mg/ml cGAMP (4 h). **(F)** RT-qPCR of fold-change in *Ifnb* expression in *LRRK2* KO vs. CT RAW 264.7 macrophages stimulated with 100 ng/ml DMXAA at indicated times. **(G)** RT-qPCR of fold-change in *Ifnb* and *Tnfa* in *LRRK2* KO vs. CT PEMs stimulated with 100 ng/ml LPS, 1 µg/ml poly(I:C), or 1 µM CL097 (4 h). (H) RT-qPCR of fold-change in *Ifnb* in *LRRK2* KO vs. CT MEFs stimulated with 1 µg/ml poly(I:C) or 100 ng/ml LPS. Data represented as means +/- S.E.M. *p<0.05, **p<0.01, ***p<0.005.

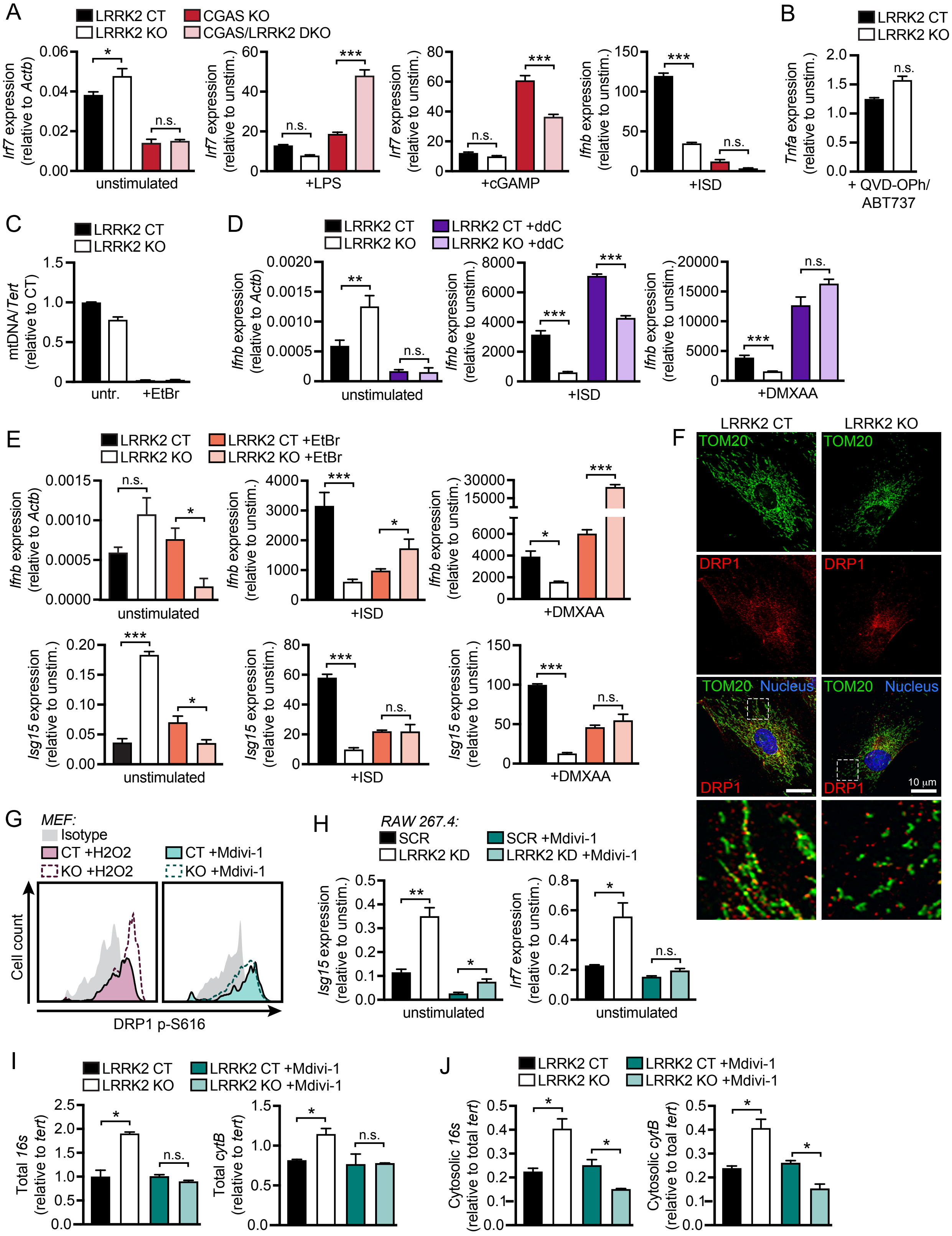


**Figure S3.** (A) RT-qPCR of *Irf7* expression normalized to *Actb* and fold-change in *Irf7* expression after stimulation with 100 ng/ml LPS or transfection with 1µg/ml cGAMP or 1µg/ml ISD (4 h) in BMDMs from CT, *LRRK2* KO, *cGAS* KO, and *LRRK2* KO/*cGAS* double KO (DKO) mice. **(B)** RT-qPCR of fold-change in *Tnfa* expression after treatment with 10 µM ABT737 and 10 µM QVD-OPh in *LRRK2* KO vs. CT BMDMs. **(C)** qPCR of ratio of *dLOOP* (mitochondrial DNA) to *Tert* (nuclear) in CT and *LRRK2* KO RAW 264.7 macrophages (normalized to CT = 1) after 4 days of 300 ng/ml ethidium bromide (EtBr) treatment. **(D)** RT-qPCR of *Ifnb* expression normalized to *Actb* and fold-change in *Isg15* expression following transfection of 1 µg/ml ISD (4 h) or 50 ng/ml DMXAA (2 h) in *LRRK2* KO vs. CT ddC-treated RAW 264.7 macrophages. **(E)** As in (D) but for Ifnb and Isg15 expression in EtBr-treated cells. (F) Immunofluorescence microscopy of total DRP1 in *LRRK2* KO vs. CT MEFs. DRP1 (red); TOM20 (green); nucleus (blue). **(G)** Histogram of counts of phospho-S616 Drp1 in *LRRK2* KO vs. CT MEFs as measured by flow cytometry after treatment with 100 µM $H_2O_2$ (1 h) or 50 µM Mdivi-1 (12 h). **(H)** RT-qPCR of *Isg15* and *Irf7* expression normalized to *Actb* in LRRK2 KD vs. SCR RAW 264.7 macrophages with or without 50 µM Mdivi-1 (12 h). **(I)** qPCR of total *16s* and *cytB* (mitochondrial DNA) relative to *tert* (nuclear DNA) in *LRRK2* KO vs. CT MEFs treated with 50 µM Mdivi-1 (12 h). **(J)** As in (I) but cytosolic *16s* and *cytB*. Data represented as means +/- S.E.M. *p<0.05, **p<0.01, ***p<0.005.

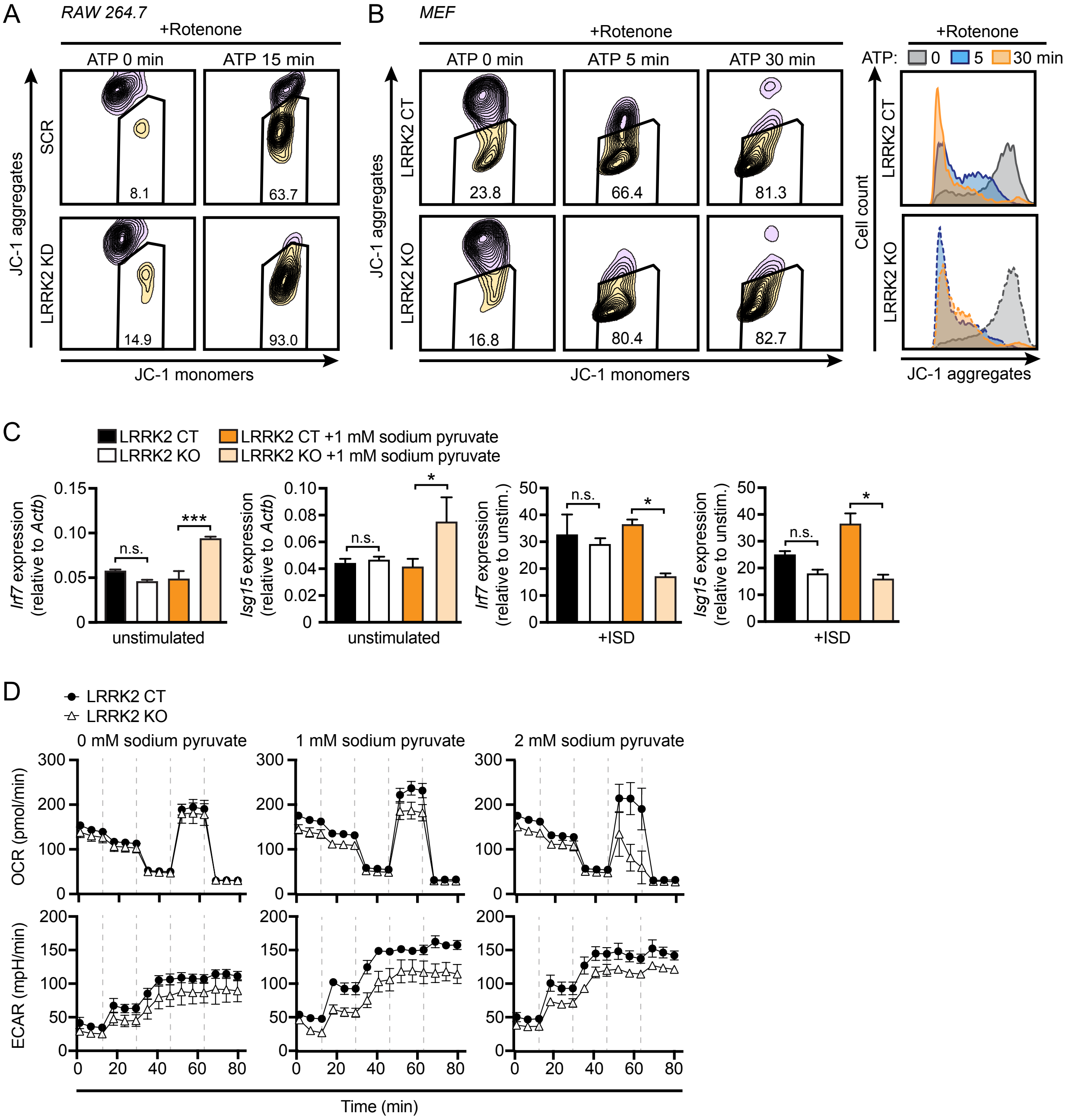


**Figure S4.** (A) Flow cytometry of JC-1 aggregates (610/20) (normal membrane potential) vs. monomers (520/50) (low membrane potential) in *LRRK2* KD vs. SCR RAW 264.7 macrophages after treatment with 2.5 µM rotenone (3 h) followed by 5 µM ATP for indicated times. **(B)** As in (A) but in MEFs. **(C)** RT-qPCR of *Irf7* and *Isg15* expression normalized to *Actb* and fold-change in *Irf7* and *Isg15* expression following transfection of 1 µg/ml ISD (4 h) in *LRRK2* KO vs. CT BMDMs grown with or without 1 mM sodium pyruvate for 24 h. **(D)** Seahorse metabolic analysis of oxygen consumption rate (OCR) and extracellular acidification rate (ECAR) in *LRRK2* KO vs. CT BMDMs grown with increasing concentrations of sodium pyruvate (0, 1, and 2 mM). Data represented as means +/- S.E.M. *p<0.05, **p<0.01, ***p<0.005.

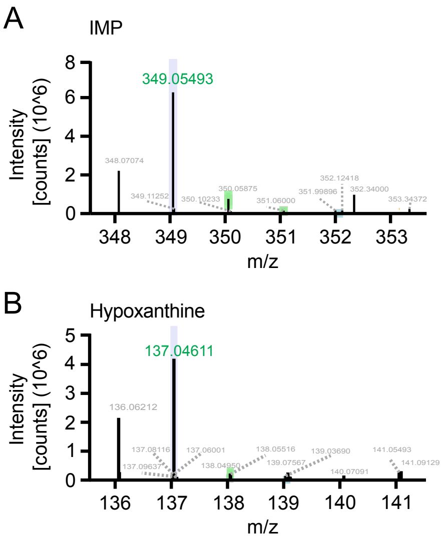


**Figure S5. (A)** LC-MS/MS analysis of *LRRK2* KO vs. CT BMDMs showing IMP. **(B)** As in (A) but for hypoxanthine.

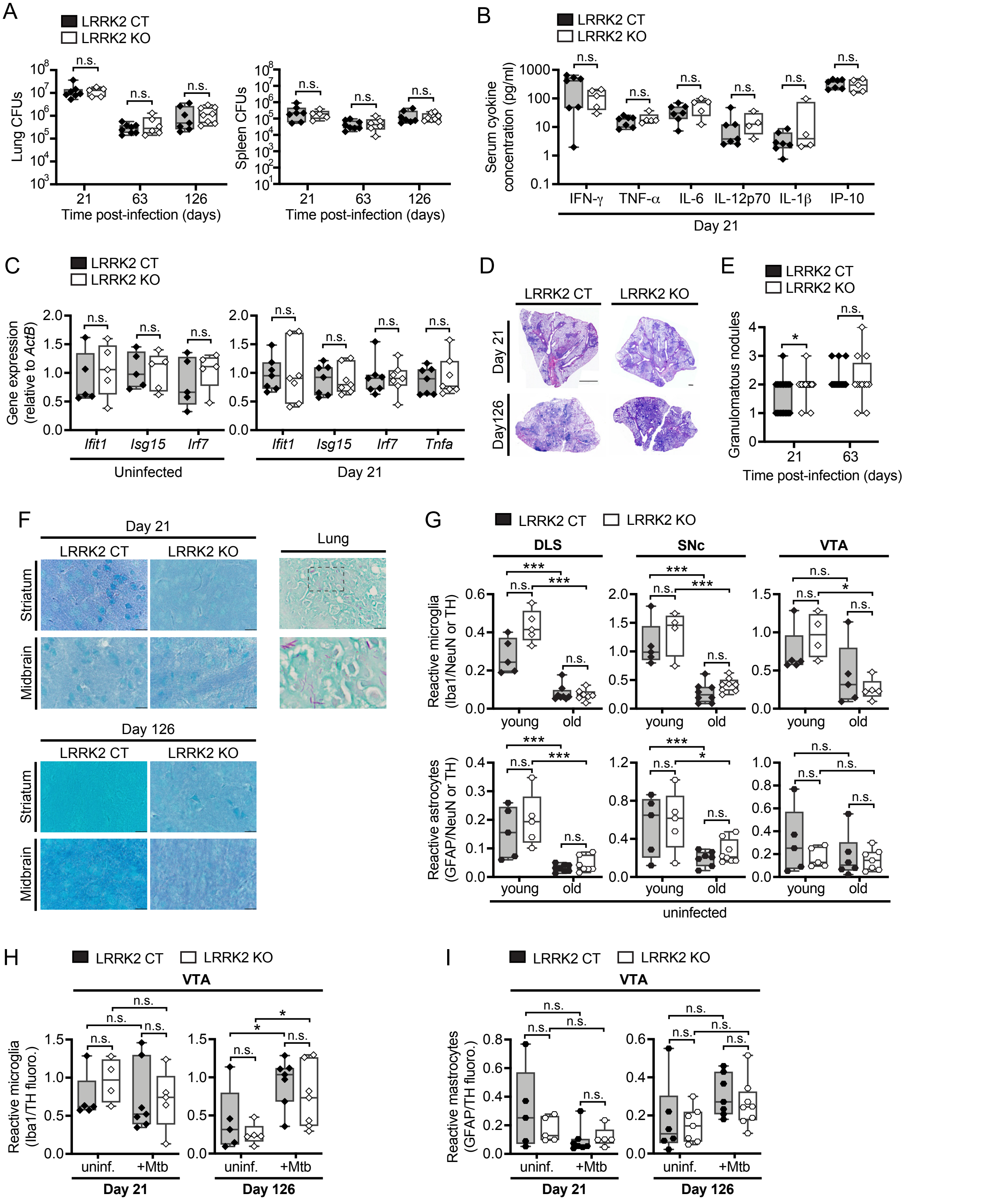


**Figure S6. (A)** Colony forming units (CFUs) recovered from lungs and spleens of Mtb-infected *LRRK2* KO vs. CT mice at 21, 63, and 126 days post-infection. **(B)** Serum cytokine concentrations in Mtb-infected *LRRK2* KO and CT mice at 21 days post-infection. **(C)** RT-qPCR of cytokine expression in lung homogenates from *LRRK2* KO vs. CT mice that were uninfected or infected with Mtb for 21 days. **(D)** H&E-stained lung sections showing inflammatory nodules in lungs of Mtb-infected *LRRK2* KO and CT mice at 21 or 126 days post-infection. **(E)** Semi-quantitative histology score of inflammatory nodules shown in (D). **(F)** Representative acid-fast stained sections of the striatum, midbrain, and lung from Mtb-infected *LRRK2* KO vs. CT mice. **(G)** Quantification of microglia and astrocyte reactivity in the DLS, SNc, and VTA as measured by Iba1 or GFAP fluorescence relative to NeuN or TH fluorescence in uninfected young and old *LRRK2* KO and CT mice. **(H)** Quantification of microglial reactivity in the VTA as measured by Iba1 fluorescence relative to TH fluorescence in *LRRK2* KO and CT mice infected with Mtb for 21 or 126 days compared to uninfected age-matched controls. **(I)** As in (H) but astrocyte reactivity measured by GFAP fluorescence. Data represented as means +/- S.E.M. *p<0.05, **p<0.01, ***p<0.005.
